## Supplementart Material for "ECOLE: Learning to call copy number variants on whole exome sequencing data"

### Supplementary Material
### for
### ECOLE: Learning to call copy number variants on whole exome sequening data

#### 1 Supplementary Tables

**Supplementary Table 1.** The performance comparison of the WES-based CNV callers on the 1000 Genomes data set (test set). CNVnator calls on the matched WGS samples are used as the ground truth. CNVkit and Control-FREEC return exact (integer) copy number predictions, which are discretized into deletion, duplication, and no-call. We also used the DECoNT tool to polish call sets of all considered tools which are denoted by DECONT-*tool_name*. Bold indicates the best result in that category.

| **TOOLS** | **DEL Precision** | **DUP Precision** | **Overall Precision** | **DEL Recall** | **DUP Recall** | **Overall Recall** | **DEL F1 Score** | **DUP F1 Score** | **Overall F1 Score** |
|---|---|---|---|---|---|---|---|---|---|
| Control-FREEC | 0.339 | 0.332 | 0.336 | 0.472 | 0.084 | 0.276 | 0.394 | 0.135 | 0.303 |
| CNVkit | 0.340 | 0.336 | 0.338 | 0.321 | 0.092 | 0.206 | 0.330 | 0.144 | 0.255 |
| XHMM | 0.407 | 0.434 | 0.421 | 0.080 | 0.071 | 0.076 | 0.134 | 0.122 | 0.129 |
| CONIFER | 0.317 | 0.105 | 0.211 | 0.004 | 0.007 | 0.006 | 0.008 | 0.013 | 0.012 |
| CODEX2 | 0.007 | 0.009 | 0.008 | 0.187 | 0.157 | 0.172 | 0.013 | 0.017 | 0.015 |
| GATK | 0.013 | 0.028 | 0.021 | 0.074 | 0.140 | 0.107 | 0.023 | 0.046 | 0.035 |
| DECoNT-Control-FREEC | 0.341 | 0.414 | 0.371 | 0.621 | 0.049 | 0.335 | 0.440 | 0.087 | 0.352 |
| DECoNT-CNVkit | 0.325 | 0.338 | 0.332 | 0.160 | 0.171 | 0.165 | 0.214 | 0.227 | 0.220 |
| DECoNT-XHMM | 0.615 | 0.652 | 0.634 | 0.086 | 0.075 | 0.081 | 0.151 | 0.135 | 0.144 |
| DECoNT-CONIFER | 0.203 | 0.152 | 0.178 | 0.011 | 0.003 | 0.007 | 0.021 | 0.006 | 0.013 |
| DECoNT-CODEX2 | 0.009 | 0.013 | 0.011 | 0.074 | 0.085 | 0.080 | 0.016 | 0.023 | 0.019 |
| ECOLE | **0.834** | **0.703** | **0.769** | **0.774** | **0.470** | **0.622** | **0.803** | **0.564** | **0.683** |

**Supplementary Table 2.** Confusion Matrices for the performance comparison based on CNVnator calls used as the ground truth for the 1000 Genomes data set (test set). These produce the precision and recall results in Supplementary Table 1.

| | | **Ground Truth** | | |
|---|---|---|---|---|
| TOOLS | **Predicted** | NO CALL | DUP | DEL |
| Control-FREEC | NO CALL | 7395972 | 6874622 | 7058879 |
| | DUP | 1324171 | 1284308 | 1248352 |
| | DEL | 7542829 | 6966242 | 7455429 |
| DECoNT-Control-FREEC | NO CALL | 5784309 | 5378215 | 5518070 |
| | DUP | 589108 | 734984 | 450704 |
| | DEL | 9889555 | 9011973 | 9793886 |
| CNVkit | NO CALL | 2047096 | 1938886 | 1921362 |
| | DUP | 299837 | 296390 | 284186 |
| | DEL | 1049869 | 974144 | 1043521 |
| DECoNT-CNVkit | NO CALL | 2290912 | 2124041 | 2205063 |
| | DUP | 557237 | 550779 | 521755 |
| | DEL | 548653 | 534600 | 522251 |
| XHMM | NO CALL | 47451741 | 417408 | 302298 |
| | DUP | 38137 | 32298 | 3916 |
| | DEL | 33510 | 4648 | 26294 |
| DECoNT-XHMM | NO CALL | 47489289 | 419118 | 303044 |
| | DUP | 17296 | 34214 | 953 |
| | DEL | 16803 | 1022 | 28511 |
| CONIFER | NO CALL | 15551274 | 174086 | 107047 |
| | DUP | 9484 | 1263 | 1280 |
| | DEL | 639 | 267 | 422 |
| DECoNT-CONIFER | NO CALL | 15554000 | 174200 | 107128 |
| | DUP | 3323 | 666 | 391 |
| | DEL | 4074 | 750 | 1230 |
| CODEX2 | NO CALL | 31430578 | 300514 | 218459 |
| | DUP | 7445915 | 71440 | 51836 |
| | DEL | 8646895 | 82400 | 62213 |
| DECoNT-CODEX2 | NO CALL | 41897336 | 395586 | 283708 |
| | DUP | 2834559 | 38655 | 24316 |
| | DEL | 2791493 | 20113 | 24484 |
| GATK | NO CALL | 27352941 | 226073 | 181310 |
| | DUP | 1369519 | 39463 | 13770 |
| | DEL | 1123082 | 17162 | 15517 |
| ECOLE | NO CALL | 29771946 | 148288 | 29050 |
| | DUP | 52583 | 133129 | 3650 |
| | DEL | 21013 | 1281 | 111866 |

**Supplementary Table 3.** Negative Predicted Value (NPV) score comparison of the WES-based CNV callers on the 1000 Genomes data set (test set). CNVnator calls on the matched WGS samples are used as the ground truth. Bold indicates the best result in that category.

| **TOOLS** | **DEL NPV Score** | **DUP NPV Score** | **Overall NPV Score** |
|---|---|---|---|
| Control-FREEC | 0.670 | 0.680 | 0.675 |
| DECoNT-Control-FREEC | 0.677 | 0.683 | 0.680 |
| CNVkit | 0.675 | 0.675 | 0.675 |
| DECoNT-CNVkit | 0.669 | 0.677 | 0.673 |
| XHMM | 0.994 | 0.991 | 0.992 |
| DECoNT-XHMM | 0.994 | 0.991 | 0.992 |
| CONIFER | 0.993 | 0.989 | 0.991 |
| DECoNT-CONIFER | 0.993 | 0.989 | 0.991 |
| CODEX2 | 0.993 | 0.991 | 0.992 |
| DECoNT-CODEX2 | 0.993 | 0.991 | 0.992 |
| GATK | 0.993 | 0.992 | 0.992 |
| ECOLE | **0.999** | **0.995** | **0.997** |

**Supplementary Table 4.** NPA (Specificity) score comparison of the WES-based CNV callers on the 1000 Genomes data set (test set). CNVnator calls on the matched WGS samples are used as the ground truth. Bold indicates the best result in that category

| **TOOLS** | **DEL NPA Score** | **DUP NPA Score** | **Overall NPA Score** |
|---|---|---|---|
| Control-FREEC | 0.538 | 0.92 | 0.729 |
| DECoNT-Control-FREEC | 0.398 | 0.968 | 0.683 |
| CNVkit | 0.694 | 0.912 | 0.803 |
| DECoNT-CNVkit | 0.836 | 0.838 | 0.837 |
| XHMM | 0.999 | 0.999 | 0.999 |
| DECoNT-XHMM | **1.0** | **1.0** | **1.0** |
| CONIFER | **1.0** | 0.999 | **1.0** |
| DECoNT-CONIFER | **1.0** | **1.0** | **1.0** |
| CODEX2 | 0.818 | 0.843 | 0.831 |
| DECoNT-CODEX2 | 0.941 | 0.94 | 0.941 |
| GATK | 0.962 | 0.954 | 0.958 |
| ECOLE | 0.999 | 0.998 | 0.999 |

**Supplementary Table 5.** Confusion Matrices for the performance comparison based on CNVnator calls used as the ground truth for the 1000 Genomes data set (28 samples obtained from S. Girirajan). These produce the precision and recall results in Table 1.

| | | **Ground Truth** | | |
|---|---|---|---|---|
| TOOLS | **Predicted** | NO CALL | DUP | DEL |
| CNLearn | NO CALL | 5324268 | 44745 | 38671 |
| | DUP | 1572 | 463 | 59 |
| | DEL | 883 | 6 | 81 |
| ECOLE | NO CALL | 5142291 | 24449 | 17584 |
| | DUP | 10369 | 23114 | 440 |
| | DEL | 3883 | 165 | 20160 |

**Supplementary Table 6.** Performance comparison on the NA12878 sample using various platforms. Bold indicates the best result in that category. CNVnator calls on the matched WGS sample are used as the semi-ground truth.

| Platform | Tool | DEL Precision | DUP Precision | Overall Precision | DEL Recall | DUP Recall | Overall Recall | DEL F1 Score | DUP F1 Score | Overall F1 Score |
|---|---|---|---|---|---|---|---|---|---|---|
| BGI 500 | Control-FREEC | 0.150 | 0.285 | 0.218 | 0.009 | 0.338 | 0.174 | 0.017 | 0.309 | 0.193 |
| | XHMM | 0.042 | 0.006 | 0.024 | 0.010 | 0.001 | 0.006 | 0.016 | 0.002 | 0.010 |
| | CONIFER | 0.002 | 0.000 | 0.001 | 0.076 | 0.000 | 0.038 | 0.004 | 0.000 | 0.002 |
| | CODEX2 | 0.007 | 0.007 | 0.007 | 0.179 | 0.101 | 0.140 | 0.013 | 0.013 | 0.013 |
| | DECoNT-Control-FREEC | 0.345 | 0.419 | 0.382 | **0.665** | 0.031 | 0.348 | 0.454 | 0.057 | 0.364 |
| | DECoNT-XHMM | 0.041 | 0.017 | 0.029 | 0.010 | 0.001 | 0.006 | 0.016 | 0.002 | 0.043 |
| | DECoNT-CONIFER | 0.010 | 0.001 | 0.006 | 0.052 | 0.010 | 0.031 | 0.017 | 0.002 | 0.010 |
| | DECoNT-CODEX2 | 0.023 | 0.026 | 0.025 | 0.097 | 0.096 | 0.097 | 0.037 | 0.041 | 0.040 |
| | ECOLE | **0.703** | **0.588** | **0.646** | 0.473 | **0.378** | **0.426** | **0.566** | **0.460** | **0.513** |
| HiSeq 4000 | Control-FREEC | 0.118 | 0.280 | 0.199 | 0.017 | 0.227 | 0.122 | 0.029 | 0.251 | 0.151 |
| | XHMM | 0.031 | 0.190 | 0.111 | 0.017 | 0.003 | 0.010 | 0.022 | 0.006 | 0.018 |
| | CONIFER | 0.035 | 0.000 | 0.018 | 0.019 | 0.000 | 0.010 | 0.025 | 0.000 | 0.013 |
| | CODEX2 | 0.008 | 0.014 | 0.011 | 0.185 | 0.213 | 0.199 | 0.015 | 0.026 | 0.021 |
| | DECoNT-Control-FREEC | 0.337 | 0.426 | 0.382 | 0.455 | 0.024 | 0.240 | 0.387 | 0.045 | 0.294 |
| | DECoNT-XHMM | 0.043 | 0.017 | 0.030 | 0.016 | 0.001 | 0.009 | 0.023 | 0.002 | 0.014 |
| | DECoNT-CONIFER | 0.018 | 0.161 | 0.090 | 0.002 | 0.090 | 0.046 | 0.004 | 0.115 | 0.061 |
| | DECoNT-CODEX2 | 0.007 | 0.007 | 0.007 | 0.034 | 0.075 | 0.055 | 0.012 | 0.013 | 0.012 |
| | ECOLE | **0.926** | **0.514** | **0.720** | **0.538** | **0.450** | **0.494** | **0.680** | **0.480** | **0.586** |
| MGISEQ 2000 | Control-FREEC | 0.150 | 0.285 | 0.218 | 0.009 | 0.338 | 0.174 | 0.017 | 0.309 | 0.193 |
| | XHMM | 0.042 | 0.006 | 0.024 | 0.010 | 0.001 | 0.006 | 0.016 | 0.002 | 0.010 |
| | CONIFER | 0.002 | 0.000 | 0.001 | 0.076 | 0.000 | 0.038 | 0.004 | 0.000 | 0.002 |
| | CODEX2 | 0.007 | 0.007 | 0.007 | 0.179 | 0.101 | 0.140 | 0.013 | 0.013 | 0.013 |
| | DECoNT-Control-FREEC | 0.345 | 0.419 | 0.382 | **0.665** | 0.031 | 0.348 | 0.454 | 0.057 | 0.364 |
| | DECoNT-XHMM | 0.041 | 0.017 | 0.029 | 0.010 | 0.001 | 0.006 | 0.016 | 0.002 | 0.043 |
| | DECoNT-CONIFER | 0.010 | 0.001 | 0.006 | 0.052 | 0.010 | 0.031 | 0.017 | 0.002 | 0.010 |
| | DECoNT-CODEX2 | 0.023 | 0.026 | 0.025 | 0.097 | 0.096 | 0.097 | 0.037 | 0.041 | 0.040 |
| | ECOLE | **0.689** | **0.587** | **0.638** | 0.472 | **0.381** | **0.427** | **0.560** | **0.462** | **0.512** |
| NovaSeq 6000 | Control-FREEC | 0.095 | 0.288 | 0.192 | 0.012 | 0.059 | 0.036 | 0.021 | 0.097 | 0.060 |
| | XHMM | 0.035 | 0.292 | 0.163 | 0.016 | 0.005 | 0.011 | 0.022 | 0.010 | 0.021 |
| | CONIFER | 0.000 | 0.000 | 0.000 | 0.000 | 0.000 | 0.000 | 0.000 | 0.000 | 0.000 |
| | CODEX2 | 0.008 | 0.018 | 0.013 | 0.228 | 0.277 | 0.253 | 0.015 | 0.034 | 0.025 |
| | DECoNT-Control-FREEC | 0.278 | 0.406 | 0.342 | 0.106 | 0.016 | 0.061 | 0.153 | 0.031 | 0.104 |
| | DECoNT-XHMM | 0.049 | 0.095 | 0.072 | 0.015 | 0.003 | 0.009 | 0.023 | 0.006 | 0.016 |
| | DECoNT-CONIFER | 0.000 | 0.000 | 0.000 | 0.000 | 0.000 | 0.000 | 0.000 | 0.000 | 0.000 |
| | DECoNT-CODEX2 | 0.016 | 0.015 | 0.016 | 0.118 | 0.181 | 0.150 | 0.028 | 0.028 | 0.029 |
| | ECOLE | **0.710** | **0.542** | **0.626** | **0.361** | **0.468** | **0.415** | **0.479** | **0.502** | **0.499** |

**Supplementary Table 7.** Confusion Matrices for the performance comparison based on CNVnator calls used as the ground truth for the NA12878 sample calls made using the BGISEQ-500 platform. These results partially produce the precision and recall results in Supplementary Table 6.

| | | **Ground Truth** | | |
|---|---|---|---|---|
| TOOLS | **Predicted** | NO CALL | DUP | DEL |
| Control-FREEC | NO CALL | 42188 | 37096 | 39088 |
| | DUP | 24112 | 20216 | 26429 |
| | DEL | 1098 | 2395 | 619 |
| DECoNT-Control-FREEC | NO CALL | 21936 | 18665 | 20979 |
| | DUP | 1408 | 1828 | 1121 |
| | DEL | 44054 | 39214 | 44036 |
| XHMM | NO CALL | 189471 | 1288 | 1763 |
| | DUP | 312 | 2 | 0 |
| | DEL | 328 | 60 | 17 |
| DECoNT-XHMM | NO CALL | 189670 | 1288 | 1763 |
| | DUP | 113 | 2 | 0 |
| | DEL | 328 | 60 | 17 |
| CONIFER | NO CALL | 125282 | 1106 | 1644 |
| | DUP | 0 | 0 | 0 |
| | DEL | 64829 | 244 | 136 |
| DECoNT-CONIFER | NO CALL | 168871 | 1165 | 1659 |
| | DUP | 12432 | 13 | 28 |
| | DEL | 8808 | 172 | 93 |
| CODEX2 | NO CALL | 128842 | 960 | 1408 |
| | DUP | 18375 | 137 | 54 |
| | DEL | 42894 | 253 | 318 |
| DECoNT-CODEX2 | NO CALL | 178118 | 1178 | 1573 |
| | DUP | 4771 | 129 | 34 |
| | DEL | 7222 | 43 | 173 |
| ECOLE | NO CALL | 189524 | 738 | 912 |
| | DUP | 332 | 511 | 26 |
| | DEL | 255 | 101 | 842 |

**Supplementary Table 8.** Confusion Matrices for the performance comparison based on CNVnator calls used as the ground truth for the NA12878 sample calls made using HiSeq 4000 platform. These results partially produce the precision and recall results in Supplementary Table 6.

| | | **Ground Truth** | | |
|---|---|---|---|---|
| TOOLS | **Predicted** | NO CALL | DUP | DEL |
| Control-FREEC | NO CALL | 48513 | 40206 | 46702 |
| | DUP | 16500 | 13543 | 18321 |
| | DEL | 2385 | 5958 | 1113 |
| DECoNT-Control-FREEC | NO CALL | 35959 | 29631 | 35210 |
| | DUP | 1096 | 1458 | 863 |
| | DEL | 30343 | 28618 | 30063 |
| XHMM | NO CALL | 189220 | 1304 | 1741 |
| | DUP | 8 | 4 | 9 |
| | DEL | 883 | 42 | 30 |
| DECoNT-XHMM | NO CALL | 189450 | 1314 | 1745 |
| | DUP | 51 | 1 | 6 |
| | DEL | 610 | 35 | 29 |
| CONIFER | NO CALL | 189323 | 1196 | 1746 |
| | DUP | 0 | 0 | 0 |
| | DEL | 788 | 154 | 34 |
| DECoNT-CONIFER | NO CALL | 189345 | 1220 | 1760 |
| | DUP | 613 | 121 | 17 |
| | DEL | 153 | 9 | 3 |
| CODEX2 | NO CALL | 128405 | 759 | 1337 |
| | DUP | 20425 | 287 | 114 |
| | DEL | 41281 | 304 | 329 |
| DECoNT-CODEX2 | NO CALL | 166852 | 1203 | 1574 |
| | DUP | 14341 | 101 | 145 |
| | DEL | 8918 | 46 | 61 |
| ECOLE | NO CALL | 189494 | 735 | 794 |
| | DUP | 548 | 608 | 28 |
| | DEL | 69 | 7 | 958 |

**Supplementary Table 9.** Confusion Matrices for the performance comparison based on CNVnator calls used as the ground truth for the NA12878 sample calls made using the MGISEQ-2000 platform. These results partially produce the precision and recall results in Supplementary Table 6.

| | | **Ground Truth** | | |
|---|---|---|---|---|
| TOOLS | **Predicted** | NO CALL | DUP | DEL |
| Control-FREEC | NO CALL | 30063 | 37096 | 39088 |
| | DUP | 24112 | 20216 | 26429 |
| | DEL | 1098 | 2395 | 619 |
| DECoNT-Control-FREEC | NO CALL | 21936 | 18665 | 20979 |
| | DUP | 1408 | 1828 | 1121 |
| | DEL | 44054 | 39214 | 44036 |
| XHMM | NO CALL | 189471 | 1288 | 1763 |
| | DUP | 312 | 2 | 0 |
| | DEL | 328 | 60 | 17 |
| DECoNT-XHMM | NO CALL | 189670 | 1288 | 1763 |
| | DUP | 113 | 2 | 0 |
| | DEL | 328 | 60 | 17 |
| CONIFER | NO CALL | 125282 | 1106 | 1644 |
| | DUP | 0 | 0 | 0 |
| | DEL | 64829 | 244 | 136 |
| DECoNT-CONIFER | NO CALL | 168871 | 1165 | 1659 |
| | DUP | 12432 | 13 | 28 |
| | DEL | 8808 | 172 | 93 |
| CODEX2 | NO CALL | 128842 | 960 | 1408 |
| | DUP | 18375 | 137 | 54 |
| | DEL | 42894 | 253 | 318 |
| DECoNT-CODEX2 | NO CALL | 178118 | 1178 | 1573 |
| | DUP | 4771 | 129 | 34 |
| | DEL | 7222 | 43 | 173 |
| ECOLE | NO CALL | 189497 | 735 | 913 |
| | DUP | 336 | 514 | 26 |
| | DEL | 278 | 101 | 841 |

**Supplementary Table 10.** Confusion Matrices for the performance comparison based on CNVnator calls used as the ground truth for the NA12878 sample calls made using NovaSeq 6000 platform. These results partially produce the precision and recall results in Supplementary Table 6.

| | | **Ground Truth** | | |
|---|---|---|---|---|
| TOOLS | **Predicted** | NO CALL | DUP | DEL |
| Control-FREEC | NO CALL | 61250 | 50528 | 60732 |
| | DUP | 4124 | 3540 | 4594 |
| | DEL | 2024 | 5639 | 810 |
| DECoNT-Control-FREEC | NO CALL | 58783 | 48391 | 58477 |
| | DUP | 761 | 952 | 631 |
| | DEL | 7854 | 10364 | 7028 |
| XHMM | NO CALL | 189382 | 1288 | 1743 |
| | DUP | 8 | 7 | 9 |
| | DEL | 721 | 55 | 28 |
| DECoNT-XHMM | NO CALL | 189595 | 1295 | 1753 |
| | DUP | 38 | 4 | 0 |
| | DEL | 478 | 51 | 27 |
| CONIFER | NO CALL | 190026 | 1346 | 1780 |
| | DUP | 0 | 0 | 0 |
| | DEL | 85 | 4 | 0 |
| DECoNT-CONIFER | NO CALL | 190026 | 1346 | 1780 |
| | DUP | 0 | 0 | 0 |
| | DEL | 85 | 4 | 0 |
| CODEX2 | NO CALL | 122523 | 702 | 1161 |
| | DUP | 20298 | 374 | 213 |
| | DEL | 47290 | 274 | 406 |
| DECoNT-CODEX2 | NO CALL | 161203 | 975 | 1375 |
| | DUP | 15728 | 245 | 195 |
| | DEL | 13180 | 130 | 210 |
| ECOLE | NO CALL | 189471 | 627 | 1073 |
| | DUP | 469 | 632 | 65 |
| | DEL | 171 | 91 | 642 |

**Supplementary Table 11.** The performance comparison of ECOLE on NA12878 samples sequenced with NimbleGen SeqCap v3 and SeqCap EZ Human Exome Library v3.0 capture kits

| **Performance Metrics** | **Capture Kits** | |
|---|---|---|
| | NimbleGen SeqCap v3 | SeqCap EZ Human Exome Library v3.0 |
| DEL Precision | 0.926 | 0.858 |
| DUP Precision | 0.514 | 0.556 |
| Overall Precision | 0.720 | 0.707 |
| DEL Recall | 0.538 | 0.495 |
| DUP Recall | 0.450 | 0.495 |
| Overall Recall | 0.494 | 0.495 |
| DEL F1 Score | 0.680 | 0.628 |
| DUP F1 Score | 0.480 | 0.524 |
| Overall F1 Score | 0.586 | 0.576 |

**Supplementary Table 12.** The performance comparison of the WES-based CNV callers on the 1000 Genomes test set which contains 157 samples. Chaisson et al.'s human expert-curated calls on the matched WGS samples are used as the ground truth [1]. We also used the DECoNT tool to polish call sets of all considered tools which are denoted by DECONT-*tool_name*. ECOLE$^{FT-EXPERT}$ corresponds to the fine-tuned version of ECOLE model. Bold indicates the best result in that category.

| **TOOLS** | **DEL Precision** | **DUP Precision** | **Overall Precision** | **DEL Recall** | **DUP Recall** | **Overall Recall** | **DEL F1 Score** | **DUP F1 Score** | **Overall F1 Score** |
|---|---|---|---|---|---|---|---|---|---|
| XHMM | 0.059 | 0.008 | 0.034 | 0.008 | 0.021 | 0.015 | 0.014 | 0.012 | 0.021 |
| CONIFER | 0.000 | **1.000** | 0.500 | 0.000 | 0.076 | 0.038 | 0.000 | 0.141 | 0.071 |
| CODEX2 | 0.023 | 0.003 | 0.013 | 0.238 | 0.138 | 0.188 | 0.042 | 0.006 | 0.024 |
| DECoNT-XHMM | 0.051 | 0.010 | 0.031 | 0.006 | 0.019 | 0.013 | 0.011 | 0.013 | 0.018 |
| DECoNT-CONIFER | 0.000 | **1.000** | 0.500 | 0.000 | 0.061 | 0.031 | 0.000 | 0.115 | 0.058 |
| DECoNT-CODEX2 | 0.022 | 0.004 | 0.013 | 0.070 | 0.087 | 0.079 | 0.033 | 0.008 | 0.022 |
| ECOLE | 0.181 | 0.070 | 0.126 | 0.050 | 0.137 | 0.094 | 0.078 | 0.093 | 0.108 |
| ECOLE$^{FT-EXPERT}$ | **0.584** | 0.789 | **0.687** | **0.444** | **0.547** | **0.496** | **0.504** | **0.646** | **0.576** |

**Supplementary Table 13.** Confusion Matrices for the performance comparison based on Chaisson et al. calls used as the ground truth [1] for the 1000 Genomes data set (test set). These produce the precision and recall results in Supplementary Table 12.

| | | **Ground Truth** | | |
|---|---|---|---|---|
| TOOLS | **Predicted** | NO CALL | DUP | DEL |
| XHMM | NO CALL | 746801 | 2556 | 14310 |
| | DUP | 7149 | 56 | 67 |
| | DEL | 1894 | 12 | 119 |
| DECoNT-XHMM | NO CALL | 749417 | 2567 | 14342 |
| | DUP | 4877 | 51 | 70 |
| | DEL | 1550 | 6 | 84 |
| CONIFER | NO CALL | 188992 | 609 | 3590 |
| | DUP | 0 | 50 | 0 |
| | DEL | 0 | 0 | 0 |
| DECoNT-CONIFER | NO CALL | 188992 | 619 | 3590 |
| | DUP | 0 | 40 | 0 |
| | DEL | 0 | 0 | 0 |
| CODEX2 | NO CALL | 481333 | 1704 | 8882 |
| | DUP | 130138 | 362 | 2161 |
| | DEL | 144373 | 558 | 3453 |
| DECoNT-CODEX2 | NO CALL | 655677 | 2266 | 12338 |
| | DUP | 54791 | 228 | 1146 |
| | DEL | 45376 | 130 | 1012 |
| ECOLE | NO CALL | 748390 | 2224 | 13165 |
| | DUP | 4197 | 359 | 604 |
| | DEL | 3257 | 41 | 727 |
| ECOLE$^{FT-EXPERT}$ | NO CALL | 751069 | 1136 | 7931 |
| | DUP | 254 | 1435 | 129 |
| | DEL | 4521 | 53 | 6436 |

**Supplementary Table 14.** Negative Predicted Value (NPV) scores comparison with other tools using Chaisson et. al labels as the ground truth [1]. Bold indicates the best result in that category.

| **TOOLS** | **DEL NPV Score** | **DUP NPV Score** | **Overall NPV Score** |
|---|---|---|---|
| XHMM | 0.981 | 0.997 | 0.989 |
| DECoNT-XHMM | 0.981 | 0.997 | 0.989 |
| CONIFER | 0.981 | 0.997 | 0.989 |
| DECoNT-CONIFER | 0.981 | 0.997 | 0.989 |
| CODEX2 | 0.982 | 0.996 | 0.989 |
| DECoNT-CODEX2 | 0.981 | 0.997 | 0.989 |
| ECOLE | 0.982 | 0.997 | 0.990 |
| ECOLE$^{FT-EXPERT}$ | **0.989** | **0.998** | **0.994** |

**Supplementary Table 15.** NPA (Specificity) scores comparison with other tools using Chaisson et. al labels as the ground truth [1]. Bold indicates the best result in that category

| **TOOLS** | **DEL NPA Score** | **DUP NPA Score** | **Overall NPA Score** |
|---|---|---|---|
| XHMM | 0.997 | 0.991 | 0.994 |
| DECoNT-XHMM | 0.998 | 0.994 | 0.996 |
| CONIFER | **1.0** | **1.0** | **1.0** |
| DECoNT-CONIFER | **1.0** | **1.0** | **1.0** |
| CODEX2 | 0.809 | 0.828 | 0.819 |
| DECoNT-CODEX2 | 0.94 | 0.927 | 0.934 |
| ECOLE | 0.996 | 0.994 | 0.995 |
| ECOLE$^{FT-EXPERT}$ | 0.994 | **1.0** | 0.997 |

**Supplementary Table 16.** Confusion Matrices for the performance comparison of calls, CNVnator calls used as the ground truth for the Somatic samples from Guo et. al. [4]. These results partially produce the precision and recall results in Table 3.

| | | **Ground Truth** | | |
|---|---|---|---|---|
| TOOLS | **Predicted** | NO CALL | DUP | DEL |
| XHMM | NO CALL | 1386707 | 951597 | 148878 |
| | DUP | 1147 | 27926 | 30 |
| | DEL | 5273 | 701 | 1886 |
| DECoNT-XHMM | NO CALL | 1387256 | 954512 | 148884 |
| | DUP | 1127 | 23602 | 178 |
| | DEL | 4744 | 2110 | 1732 |
| ECOLE | NO CALL | 1385260 | 968078 | 147230 |
| | DUP | 4561 | 10608 | 670 |
| | DEL | 3306 | 1538 | 2894 |
| ECOLE$^{FT-SOMATIC}$ | NO CALL | 908337 | 585915 | 92415 |
| | DUP | 439066 | 372057 | 37171 |
| | DEL | 45724 | 22252 | 21208 |

**Supplementary Table 17.** The performance comparison of the WES-based CNV callers on the 1000 Genomes data set (test set) on CNV segments (merged exons). CNVnator calls on the matched WGS samples are used as the ground truth. CNVkit and Control-FREEC return exact (integer) copy number predictions, which are discretized into deletion, duplication, and no-call. We also used the DECoNT tool to polish call sets of all considered tools which are denoted by DECONT-*tool_name*. See Supplementary Table 18 for corresponding confusion matrices.

| **TOOLS** | **CNV Segment Del Precision** | **CNV Segment Dup Precision** | **Overall Precision** |
|---|---|---|---|
| Control-FREEC | 0.673 | 0.107 | 0.391 |
| CNVkit | 0.590 | 0.153 | 0.371 |
| XHMM | 0.434 | 0.455 | 0.445 |
| CONIFER | 0.404 | 0.205 | 0.305 |
| CODEX2 | 0.191 | 0.044 | 0.118 |
| DECoNT-Control-FREEC | 0.661 | 0.418 | 0.540 |
| DECoNT-CNVkit | 0.604 | 0.584 | 0.594 |
| DECoNT-XHMM | **0.757** | 0.688 | 0.722 |
| DECoNT-CONIFER | 0.525 | 0.367 | 0.446 |
| DECoNT-CODEX2 | 0.356 | 0.115 | 0.236 |
| ECOLE | 0.724 | **0.721** | **0.723** |

**Supplementary Table 18.** Confusion Matrices for the performance comparison of calls on CNV segments and CNVnator calls are grouped together (CNV segments) to be used as ground truth for the 1000 Genomes data set (test set). These produce the precision results in Supplementary Table 17.

| | | **Ground Truth** | | |
|---|---|---|---|---|
| TOOLS | **Predicted** | NO CALL | DUP | DEL |
| Control-FREEC | NO CALL | 1800 | 562 | 3585 |
| | DUP | 26154 | 2787 | 4986 |
| | DEL | 7962 | 2664 | 13788 |
| DECoNT-Control-FREEC | NO CALL | 0 | 0 | 0 |
| | DUP | 9770 | 1740 | 2783 |
| | DEL | 26146 | 4273 | 19576 |
| CNVkit | NO CALL | 83 | 157 | 1330 |
| | DUP | 2967 | 589 | 1791 |
| | DEL | 2589 | 421 | 2152 |
| DECoNT-CNVkit | NO CALL | 0 | 0 | 0 |
| | DUP | 66 | 35 | 122 |
| | DEL | 188 | 24 | 115 |
| XHMM | NO CALL | 0 | 0 | 0 |
| | DUP | 5428 | 5042 | 607 |
| | DEL | 4303 | 1473 | 4438 |
| DECoNT-XHMM | NO CALL | 6421 | 887 | 676 |
| | DUP | 2255 | 5338 | 161 |
| | DEL | 1055 | 290 | 4208 |
| CONIFER | NO CALL | 0 | 0 | 0 |
| | DUP | 292 | 111 | 137 |
| | DEL | 9 | 22 | 21 |
| DECoNT-CONIFER | NO CALL | 162 | 14 | 13 |
| | DUP | 89 | 76 | 42 |
| | DEL | 50 | 43 | 103 |
| CODEX2 | NO CALL | 0 | 0 | 0 |
| | DUP | 119576 | 6655 | 24667 |
| | DEL | 95429 | 5868 | 23934 |
| DECoNT-CODEX2 | NO CALL | 164350 | 2826 | 11550 |
| | DUP | 26112 | 4837 | 10943 |
| | DEL | 22606 | 2525 | 13909 |
| ECOLE | NO CALL | 9253 | 9003 | 21057 |
| | DUP | 5919 | 17124 | 719 |
| | DEL | 4352 | 250 | 11534 |

**Supplementary Table 19.** Performance comparison with CNLearn on 1000 Genome data when using CNV segments. The results of the 28 samples for which CNLearn calls are obtained via personal communication are shown. Bold indicates the best result in that category. See Supplementary Table 20 for corresponding confusion matrices.

| **TOOLS** | **CNV Segment Del Precision** | **CNV Segment Dup Precision** | **Overall Precision** |
|---|---|---|---|
| CNLearn | 0.171 | 0.246 | 0.209 |
| ECOLE | **0.664** | **0.712** | **0.688** |

**Supplementary Table 20.** Confusion Matrices for the performance comparison of calls on CNV segments and CNVnator calls are grouped together to be used as ground truth for the 1000 Genomes data set (28 samples obtained from S. Girirajan). These produce the precision and recall results in Supplementary Table 19.

| | | **Predicted** | | |
|---|---|---|---|---|
| TOOLS | **Actual** | NO CALL | DUP | DEL |
| CNLearn | NO CALL | 0 | 0 | 0 |
| | DUP | 92 | 35 | 22 |
| | DEL | 184 | 2 | 38 |
| ECOLE | NO CALL | 1371 | 1952 | 4054 |
| | DUP | 1033 | 2985 | 238 |
| | DEL | 892 | 89 | 1953 |

**Supplementary Table 21.** Performance comparison with other tools using Chaisson et. al labels as the ground truth [1]. Results for 4 samples from this study which are left out for testing are shown. This table considers grouped CNV calls (CNV segments). Bold indicates the best result in that category. See Supplementary Table 22 for corresponding confusion matrices.

| **TOOLS** | **CNV Segment DEL Precision** | **CNV Segment DUP Precision** | **Overall Precision** |
|---|---|---|---|
| XHMM | 0.225 | 0.054 | 0.140 |
| CONIFER | 0.000 | **1.000** | 0.500 |
| CODEX2 | 0.409 | 0.019 | 0.214 |
| DECoNT-XHMM | 0.354 | 0.052 | 0.203 |
| DECoNT-CONIFER | 0.000 | **1.000** | 0.500 |
| DECoNT-CODEX2 | **0.664** | 0.033 | 0.349 |
| ECOLE | 0.425 | 0.148 | 0.286 |
| ECOLE$^{FT-EXPERT}$ | 0.487 | 0.799 | **0.643** |

**Supplementary Table 22.** Confusion Matrices for the performance comparison of calls on CNV segments and Chaisson et. al. calls used as the ground truth [1] for the 1000 Genomes data set (test set). These produce the precision results in Supplementary Table 21.

| | | **Ground Truth** | | |
|---|---|---|---|---|
| TOOLS | **Predicted** | NO CALL | DUP | DEL |
| XHMM | NO CALL | 0 | 0 | 0 |
| | DUP | 169 | 14 | 75 |
| | DEL | 108 | 9 | 34 |
| DECoNT-XHMM | NO CALL | 100 | 9 | 25 |
| | DUP | 128 | 10 | 55 |
| | DEL | 49 | 4 | 29 |
| CONIFER | NO CALL | 0 | 0 | 0 |
| | DUP | 0 | 6 | 0 |
| | DEL | 0 | 0 | 0 |
| DECoNT-CONIFER | NO CALL | 0 | 0 | 0 |
| | DUP | 1 | 2 | 0 |
| | DEL | 0 | 0 | 0 |
| CODEX2 | NO CALL | 0 | 0 | 0 |
| | DUP | 1605 | 50 | 980 |
| | DEL | 1172 | 43 | 844 |
| DECoNT-CODEX2 | NO CALL | 2263 | 51 | 597 |
| | DUP | 319 | 30 | 568 |
| | DEL | 188 | 11 | 393 |
| ECOLE | NO CALL | 347 | 55 | 634 |
| | DUP | 474 | 95 | 107 |
| | DEL | 194 | 9 | 143 |
| ECOLE$^{FT-EXPERT}$ | NO CALL | 1245 | 50 | 2128 |
| | DUP | 57 | 300 | 28 |
| | DEL | 1532 | 11 | 1463 |

**Supplementary Table 23.** Performance comparison on the NA12878 sample on CNV segments (merged exons). CNVnator calls on the matched WGS samples are used as the ground truth. Bold indicates the best result in that category. See Supplementary Tables 24-27 for corresponding confusion matrices.

| **Platform** | **Tool** | **CNV Segment Del Precision** | **CNV Segment Dup Precision** | **CNV Segment Overall Precision** |
|---|---|---|---|---|
| BGI 500 | Control-FREEC | 0.405 | 0.156 | 0.280 |
| | XHMM | 0.158 | 0.045 | 0.102 |
| | CONIFER | 0.051 | 0.000 | 0.026 |
| | CODEX2 | 0.216 | 0.040 | 0.128 |
| | DECoNT-Control-FREEC | 0.537 | 0.194 | 0.366 |
| | DECoNT-XHMM | 0.176 | 0.076 | 0.126 |
| | DECoNT-CONIFER | 0.096 | 0.010 | 0.053 |
| | DECoNT-CODEX2 | 0.436 | 0.133 | 0.285 |
| | ECOLE | **0.625** | **0.713** | **0.669** |
| HiSeq 4000 | Control-FREEC | 0.328 | 0.171 | 0.250 |
| | XHMM | 0.095 | 0.500 | 0.298 |
| | CONIFER | 0.173 | 0.000 | 0.087 |
| | CODEX2 | 0.180 | 0.030 | 0.105 |
| | DECoNT-Control-FREEC | 0.418 | 0.180 | 0.300 |
| | DECoNT-XHMM | 0.158 | 0.000 | 0.079 |
| | DECoNT-CONIFER | 0.125 | 0.500 | 0.313 |
| | DECoNT-CODEX2 | 0.331 | 0.067 | 0.200 |
| | ECOLE | **0.810** | **0.508** | **0.659** |
| MGISEQ 2000 | Control-FREEC | 0.405 | 0.156 | 0.280 |
| | XHMM | 0.158 | 0.045 | 0.102 |
| | CONIFER | 0.051 | 0.000 | 0.026 |
| | CODEX2 | 0.216 | 0.040 | 0.128 |
| | DECoNT-Control-FREEC | 0.537 | 0.194 | 0.366 |
| | DECoNT-XHMM | 0.176 | 0.076 | 0.126 |
| | DECoNT-CONIFER | 0.096 | 0.010 | 0.053 |
| | DECoNT-CODEX2 | 0.436 | 0.133 | 0.285 |
| | ECOLE | **0.615** | **0.682** | **0.648** |
| NovaSeq 6000 | Control-FREEC | 0.333 | 0.195 | 0.264 |
| | XHMM | 0.081 | **0.666** | 0.374 |
| | CONIFER | 0.000 | 0.000 | 0.000 |
| | CODEX2 | 0.189 | 0.043 | 0.116 |
| | DECoNT-Control-FREEC | 0.362 | 0.226 | 0.294 |
| | DECoNT-XHMM | 0.115 | 0.200 | 0.158 |
| | DECoNT-CONIFER | 0.000 | 0.000 | 0.000 |
| | DECoNT-CODEX2 | 0.308 | 0.135 | 0.222 |
| | ECOLE | **0.708** | 0.593 | **0.651** |

**Supplementary Table 24.** Confusion Matrices for the performance comparison of calls on CNV segments and CNVnator calls used as the ground truth for the NA12878 sample calls made using the BGISEQ-500 platform. These results partially produce the precision results in Supplementary Table 23.

| | | **Ground Truth** | | |
|---|---|---|---|---|
| TOOLS | **Predicted** | NO CALL | DUP | DEL |
| Control-FREEC | NO CALL | 16 | 9 | 71 |
| | DUP | 49 | 23 | 80 |
| | DEL | 50 | 19 | 50 |
| DECoNT-Control-FREEC | NO CALL | 0 | 0 | 0 |
| | DUP | 9 | 6 | 19 |
| | DEL | 106 | 45 | 182 |
| XHMM | NO CALL | 0 | 0 | 0 |
| | DUP | 21 | 1 | 0 |
| | DEL | 10 | 6 | 3 |
| DECoNT-XHMM | NO CALL | 10 | 1 | 0 |
| | DUP | 12 | 1 | 0 |
| | DEL | 9 | 5 | 3 |
| CONIFER | NO CALL | 0 | 0 | 0 |
| | DUP | 0 | 0 | 0 |
| | DEL | 5301 | 58 | 308 |
| DECoNT-CONIFER | NO CALL | 7076 | 48 | 280 |
| | DUP | 2022 | 24 | 152 |
| | DEL | 1504 | 44 | 184 |
| CODEX2 | NO CALL | 0 | 0 | 0 |
| | DUP | 640 | 32 | 136 |
| | DEL | 500 | 22 | 148 |
| DECoNT-CODEX2 | NO CALL | 478 | 9 | 35 |
| | DUP | 31 | 8 | 24 |
| | DEL | 55 | 3 | 45 |
| ECOLE | NO CALL | 48 | 30 | 114 |
| | DUP | 20 | 62 | 6 |
| | DEL | 36 | 0 | 60 |

**Supplementary Table 25.** Confusion Matrices for the performance comparison of calls on CNV segments and CNVnator calls used as the ground truth for the NA12878 sample calls made using HiSeq 4000 platform. These results partially produce the precision results in Supplementary Table 23.

| | | **Ground Truth** | | |
|---|---|---|---|---|
| TOOLS | **Predicted** | NO CALL | DUP | DEL |
| Control-FREEC | NO CALL | 22 | 6 | 49 |
| | DUP | 27 | 14 | 44 |
| | DEL | 125 | 53 | 97 |
| DECoNT-Control-FREEC | NO CALL | 0 | 0 | 0 |
| | DUP | 17 | 7 | 15 |
| | DEL | 157 | 66 | 175 |
| XHMM | NO CALL | 0 | 0 | 0 |
| | DUP | 0 | 1 | 1 |
| | DEL | 79 | 16 | 12 |
| DECoNT-XHMM | NO CALL | 27 | 10 | 0 |
| | DUP | 6 | 0 | 2 |
| | DEL | 46 | 7 | 11 |
| CONIFER | NO CALL | 0 | 0 | 0 |
| | DUP | 0 | 0 | 0 |
| | DEL | 30 | 13 | 10 |
| DECoNT-CONIFER | NO CALL | 34 | 12 | 10 |
| | DUP | 4 | 8 | 6 |
| | DEL | 22 | 6 | 4 |
| CODEX2 | NO CALL | 0 | 0 | 0 |
| | DUP | 1006 | 38 | 214 |
| | DEL | 754 | 38 | 184 |
| DECoNT-CODEX2 | NO CALL | 607 | 9 | 54 |
| | DUP | 163 | 15 | 49 |
| | DEL | 104 | 7 | 58 |
| ECOLE | NO CALL | 76 | 45 | 151 |
| | DUP | 82 | 91 | 9 |
| | DEL | 15 | 1 | 68 |

**Supplementary Table 26.** Confusion Matrices for the performance comparison of calls on CNV segments and CNVnator calls used as the ground truth for the NA12878 sample calls made using the BGISEQ-500 platform. These results partially produce the precision results in Supplementary Table 23.

| | | **Ground Truth** | | |
|---|---|---|---|---|
| TOOLS | **Predicted** | NO CALL | DUP | DEL |
| Control-FREEC | NO CALL | 16 | 9 | 71 |
| | DUP | 49 | 23 | 80 |
| | DEL | 50 | 19 | 50 |
| DECoNT-Control-FREEC | NO CALL | 0 | 0 | 0 |
| | DUP | 9 | 6 | 19 |
| | DEL | 106 | 45 | 182 |
| XHMM | NO CALL | 0 | 0 | 0 |
| | DUP | 21 | 1 | 0 |
| | DEL | 10 | 6 | 3 |
| DECoNT-XHMM | NO CALL | 10 | 1 | 0 |
| | DUP | 12 | 1 | 0 |
| | DEL | 9 | 5 | 3 |
| CONIFER | NO CALL | 0 | 0 | 0 |
| | DUP | 0 | 0 | 0 |
| | DEL | 5301 | 58 | 308 |
| DECoNT-CONIFER | NO CALL | 7076 | 48 | 280 |
| | DUP | 2022 | 24 | 152 |
| | DEL | 1504 | 44 | 184 |
| CODEX2 | NO CALL | 0 | 0 | 0 |
| | DUP | 640 | 32 | 136 |
| | DEL | 500 | 22 | 148 |
| DECoNT-CODEX2 | NO CALL | 478 | 9 | 35 |
| | DUP | 31 | 8 | 24 |
| | DEL | 55 | 3 | 45 |
| ECOLE | NO CALL | 53 | 27 | 113 |
| | DUP | 23 | 60 | 6 |
| | DEL | 37 | 0 | 59 |

**Supplementary Table 27.** Confusion Matrices for the performance comparison of calls on CNV segments and CNVnator calls used as the ground truth for the NA12878 sample calls made using NovaSeq 6000 platform. These results partially produce the precision results in Supplementary Table 23.

| | | **Ground Truth** | | |
|---|---|---|---|---|
| TOOLS | **Predicted** | NO CALL | DUP | DEL |
| Control-FREEC | NO CALL | 15 | 4 | 26 |
| | DUP | 19 | 8 | 19 |
| | DEL | 103 | 49 | 86 |
| DECoNT-Control-FREEC | NO CALL | 0 | 0 | 0 |
| | DUP | 11 | 7 | 15 |
| | DEL | 126 | 54 | 116 |
| XHMM | NO CALL | 0 | 0 | 0 |
| | DUP | 0 | 2 | 1 |
| | DEL | 65 | 14 | 10 |
| DECoNT-XHMM | NO CALL | 24 | 6 | 2 |
| | DUP | 4 | 1 | 1 |
| | DEL | 37 | 9 | 8 |
| CONIFER | NO CALL | 0 | 0 | 0 |
| | DUP | 0 | 0 | 0 |
| | DEL | 8 | 2 | 0 |
| DECoNT-CONIFER | NO CALL | 0 | 0 | 0 |
| | DUP | 0 | 0 | 0 |
| | DEL | 16 | 4 | 0 |
| CODEX2 | NO CALL | 0 | 0 | 0 |
| | DUP | 1226 | 64 | 196 |
| | DEL | 884 | 66 | 236 |
| DECoNT-CODEX2 | NO CALL | 840 | 14 | 43 |
| | DUP | 104 | 26 | 68 |
| | DEL | 107 | 12 | 57 |
| ECOLE | NO CALL | 60 | 42 | 141 |
| | DUP | 50 | 83 | 8 |
| | DEL | 27 | 1 | 68 |

**Supplementary Table 28.** Lists of samples used for training and testing ECOLE in different scenarios are given. This data is provided as a separate Excel file.

**Supplementary Table 29.** Performance comparison with the SVM baseline model using the test set from 1000 Genome data. We use the exon-level semi-ground truth labels obtained using CNVnator. Bold indicates the best result in that category.

| **TOOLS** | **DEL Precision** | **DEL Recall** | **DUP Precision** | **DUP Recall** | **Overall Precision** | **Overall Recall** |
|---|---|---|---|---|---|---|
| SVM | 0.042 | 0.627 | 0.017 | **0.543** | 0.029 | 0.585 |
| XGBoost | 0.728 | 0.0208 | 0.561 | 0.0186 | 0.644 | 0.0197 |
| ECOLE | **0.834** | **0.774** | **0.703** | 0.470 | **0.769** | **0.622** |

**Supplementary Table 30.** The performance comparison of ECOLE on chromosome 21 and 10, against convolutional neural network (CNN)-based models that are trained and tested on chromosome 21 and 10.

| Model | DEL Precision | DUP Precision | Overall Precision | DEL Recall | DUP Recall | Overall Recall | DEL F1 Score | DUP F1 Score | Overall F1 Score |
|---|---|---|---|---|---|---|---|---|---|
| Chromosome-Specific CNN (chr 21) | 0.466 | 0.711 | 0.588 | 0.0376 | 0.00144 | 0.0195 | 0.0696 | 0.00287 | 0.0362 |
| ECOLE (only chr 21) | 0.760 | 0.950 | 0.855 | 0.959 | 0.893 | 0.926 | 0.848 | 0.920 | 0.884 |
| Chromosome-Specific CNN (chr 10) | 0.0 | 0.0 | 0.0 | 0.0 | 0.0 | 0.0 | 0.0 | 0.0 | 0.0 |
| ECOLE (only chr 10) | 0.0 | 0.0753 | 0.0377 | 0.0 | 0.585 | 0.292 | 0.0 | 0.133 | 0.0667 |

**Supplementary Table 31.** Performance comparison with Base Transformer using the test set from 1000 Genome data. We use the exon-level semi-ground truth labels obtained using CNVnator. Bold indicates the best result in that category.

| **TOOLS** | **DEL Precision** | **DEL Recall** | **DUP Precision** | **DUP Recall** | **Overall Precision** | **Overall Recall** |
|---|---|---|---|---|---|---|
| Transformer base | 0.625 | 0.00007 | 0.312 | 0.007 | 0.469 | 0.004 |
| ECOLE | **0.834** | **0.774** | **0.703** | **0.470** | **0.769** | **0.622** |

#### 2 Supplementary Figures

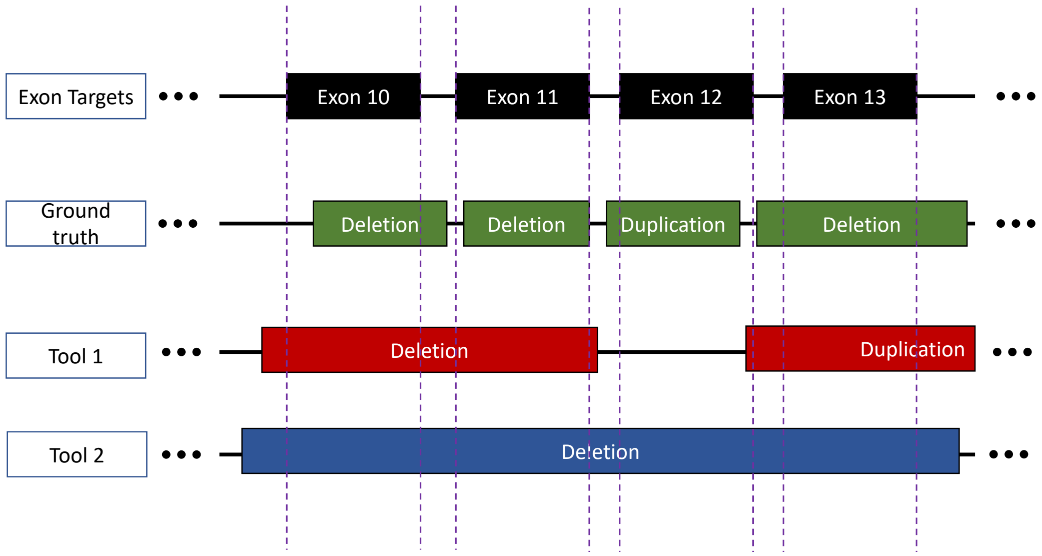


**Supplementary Figure 1.** The ground truths of exon-level CNV calling, introduced in Section 2.2, are obtained by intersecting the CNV call segments with the exons. Moreover, the CNV calls made by the tools are also obtained by intersecting them with the exons, thus we have fair comparison between different tools. If the tool or the ground truth did not report a CNV call in the target exon, then there is a no-call.In this case, we have a unique ground truth. In this example, Tool 1 has 2 TP calls for exons 10 and 11. It has 1 FP call for exon 13. Tool 2 has 3 TP calls for exons 10, 11, and 13. It has 1 FP call for exon 12. See Supplementary Figure 9 for the alternative approach.

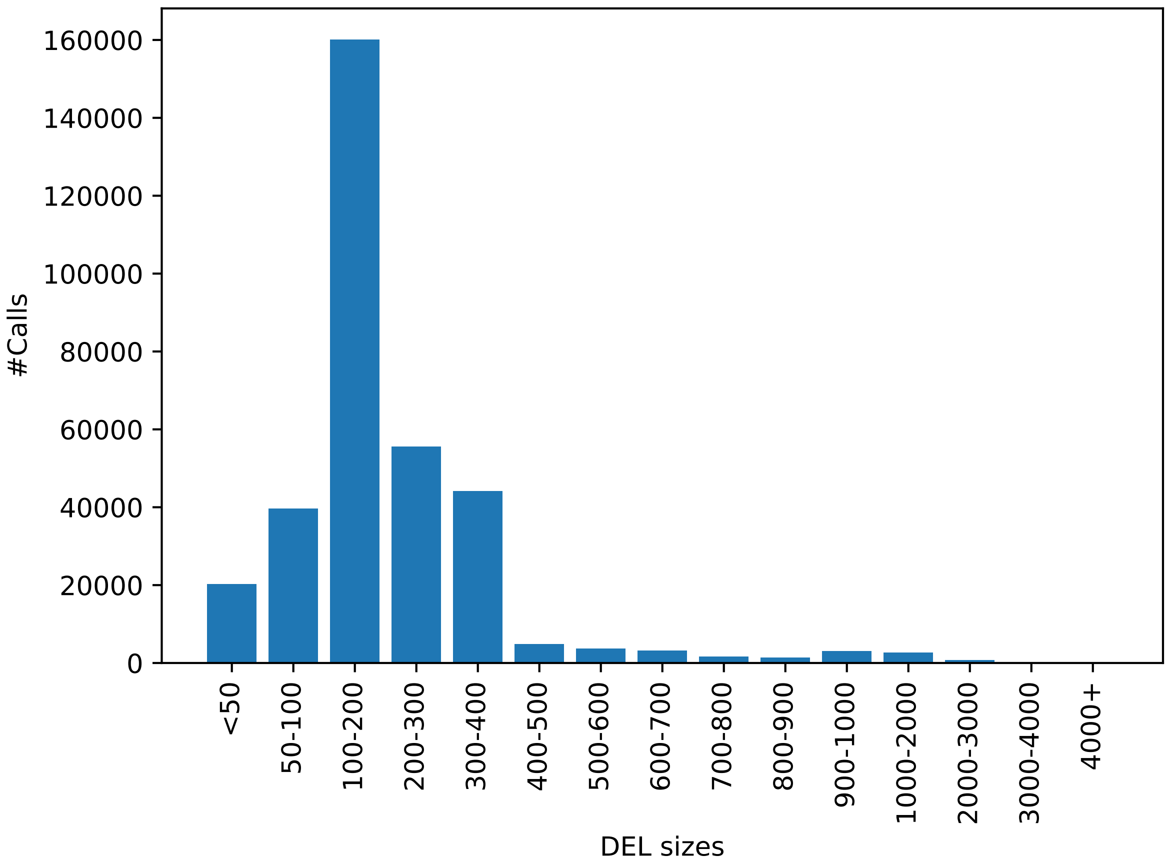


**Supplementary Figure 2.** Distribution of deletion call sizes in CNVnator predictions on the (WGS) 1000 Genomes data set (training set).

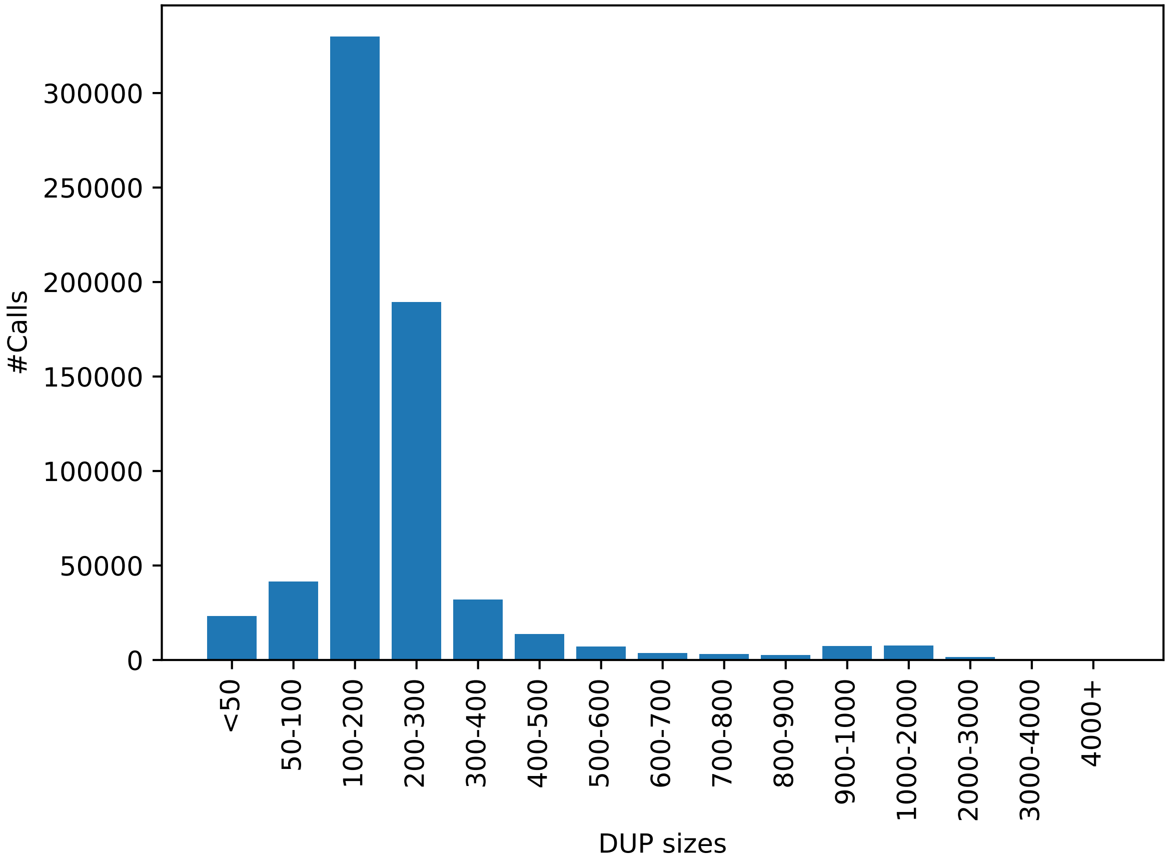


**Supplementary Figure 3.** Distribution of duplication call sizes in CNVnator predictions on the (WGS) 1000 Genomes data set (training set).

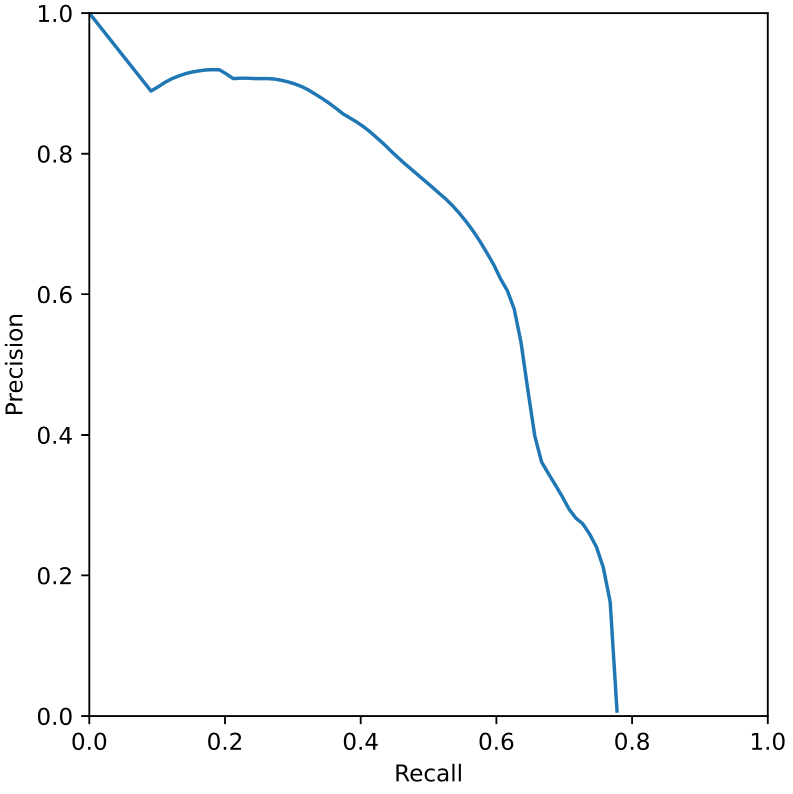


**Supplementary Figure 4.** The macro average precision-recall curve of ECOLE on 1000 Genomes data set (test set).The AUPR score of the model is 0.579

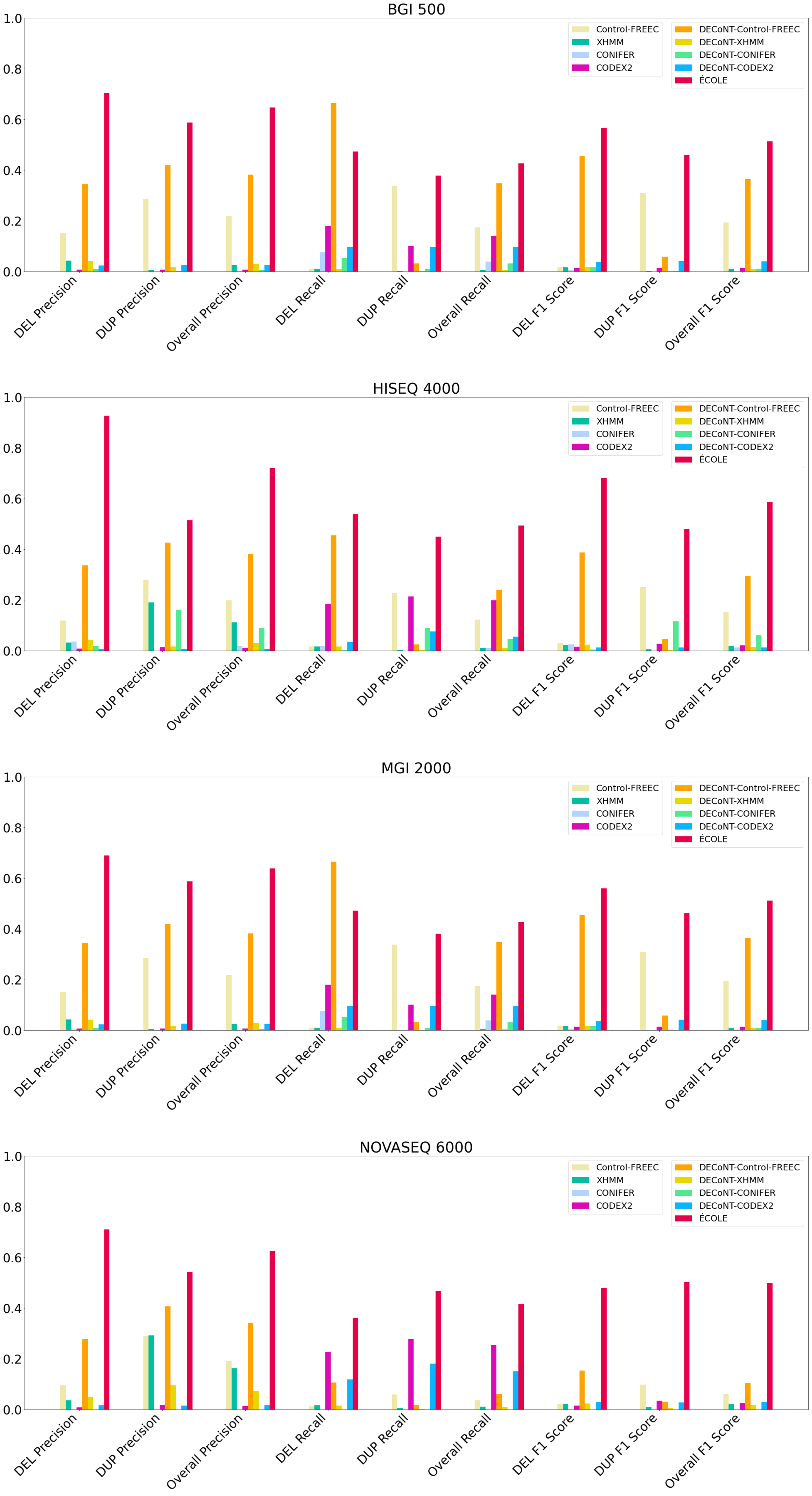


**Supplementary Figure 5.** Performance comparison with other tools on NA12878 sample. See Supplementary Table 6 for the values used to produce this plot and see Supplementary Tables 7-10 for the corresponding confusion matrices.

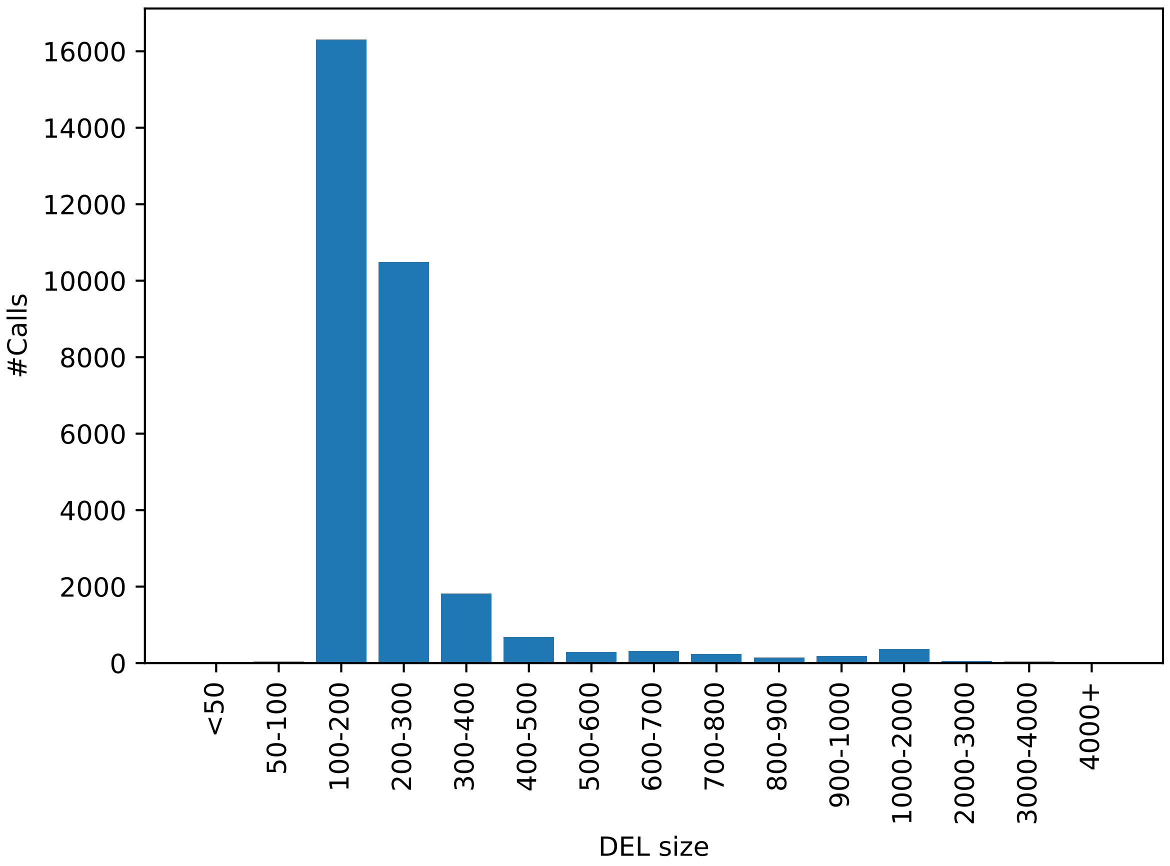


**Supplementary Figure 6.** Distribution of deletion ground-truth call sizes in Chaisson et. al. data set (test set).

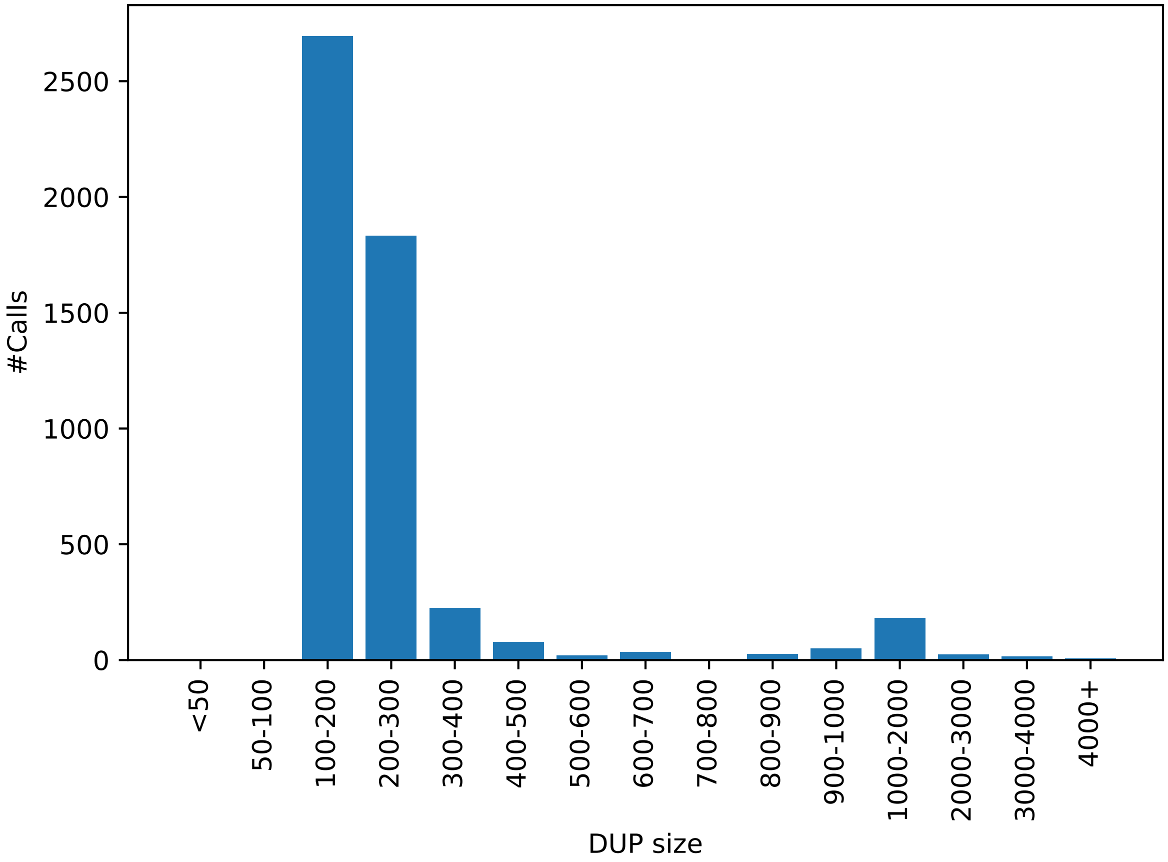


**Supplementary Figure 7.** Distribution of duplication ground-truth call sizes in Chaisson et. al. data set (test set).

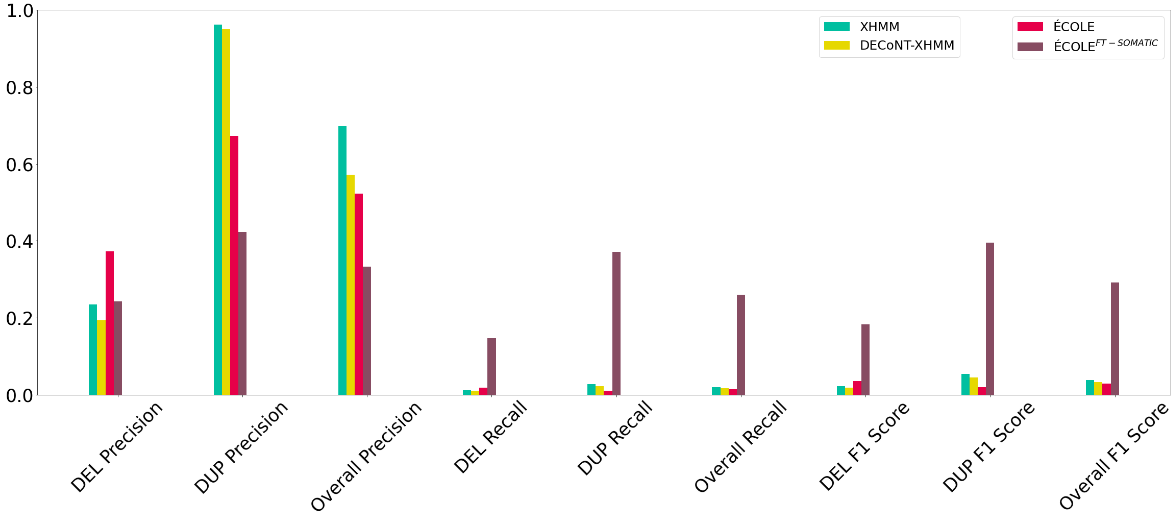


**Supplementary Figure 8.** The performance comparison with other tools on samples from Guo et. al. [4]. See Table 3 for the values used to produce this plot and see Supplementary Table 16 for the corresponding confusion matrix.

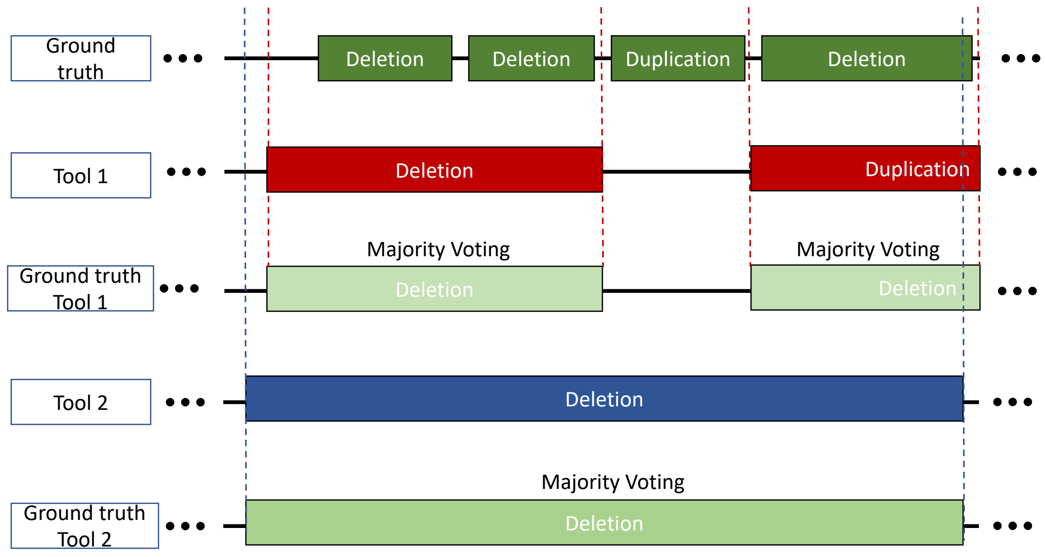


**Supplementary Figure 9.** The ground truths of merged CNV segment comparison, introduced in Section 2.3, are obtained by applying majority voting on the ground truth calls inside the target region of a CNV call made by a tool. Note that the ground truth labels are always shorter in length so majority voting is necessary to obtain the respective ground truth for a merged CNV call made by a tool. In this case, we have multiple ground truths for each tool, which prohibit the calculation of the recall fairly. Here, Tool 1 has 1 TP deletion call as the first two exons are grouped w.r.t. the calls of Tool 1, and a consensus ground truth is assigned. It has a single FP call for the last exon. Tool 2 has 1 TP deletion call as all exons are grouped due to the large call of Tool 2 which spans all exons. The consensus is that these exons have a deletion label w.r.t. majority voting.

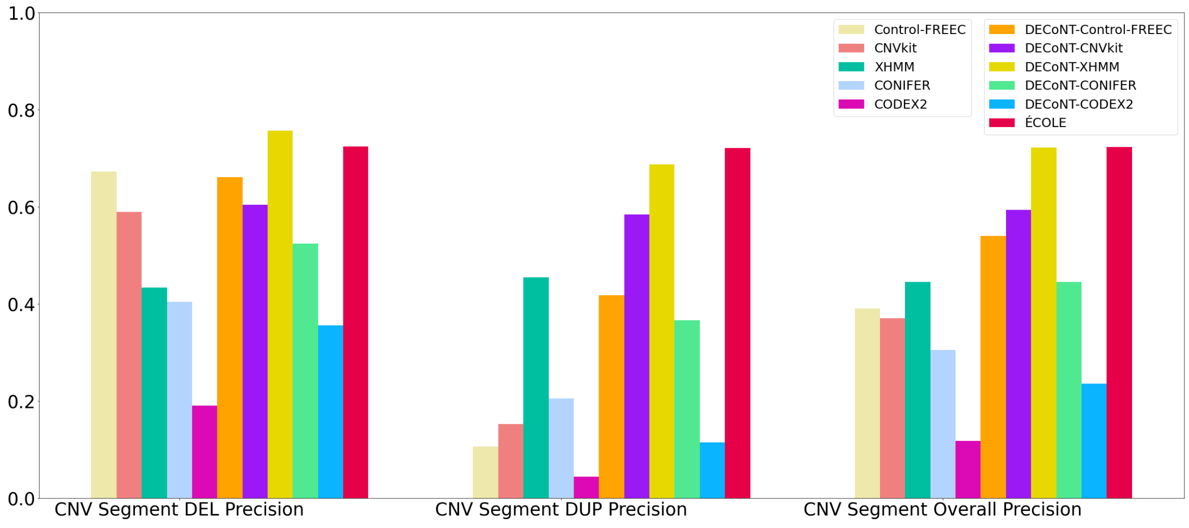


**Supplementary Figure 10.** The performance comparison of the WES-based CNV callers on the 1000 Genomes data set (test set) on CNV segments. CNVnator calls on the matched WGS samples are used as the ground truth. CNVkit and Control-FREEC return exact (integer) copy number predictions, which are discretized into deletion, duplication, and no-call. We also used the DECoNT tool to polish call sets of all considered tools which are denoted by DECONT-*tool_name*

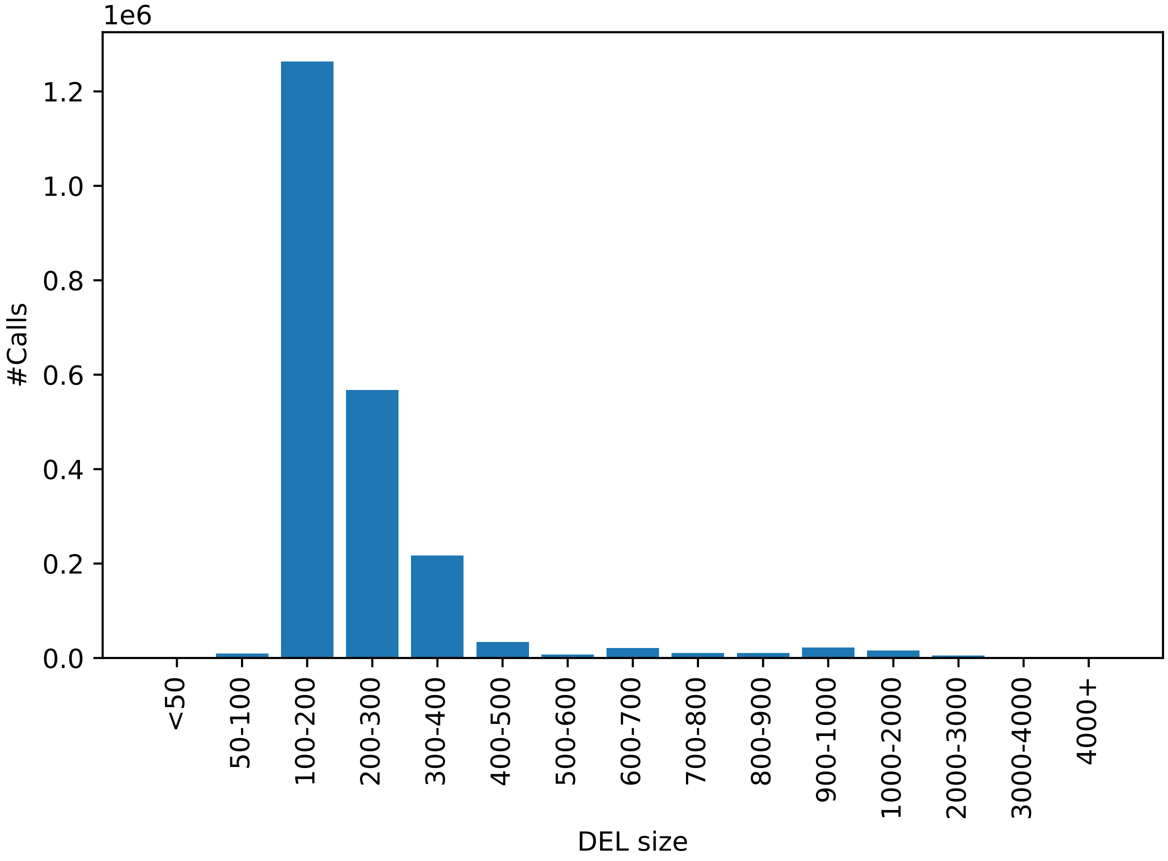


**Supplementary Figure 11.** Distribution of deletion call (merged CNV segments) sizes in CNVnator predictions on the (WGS) 1000 Genomes data set (training set).

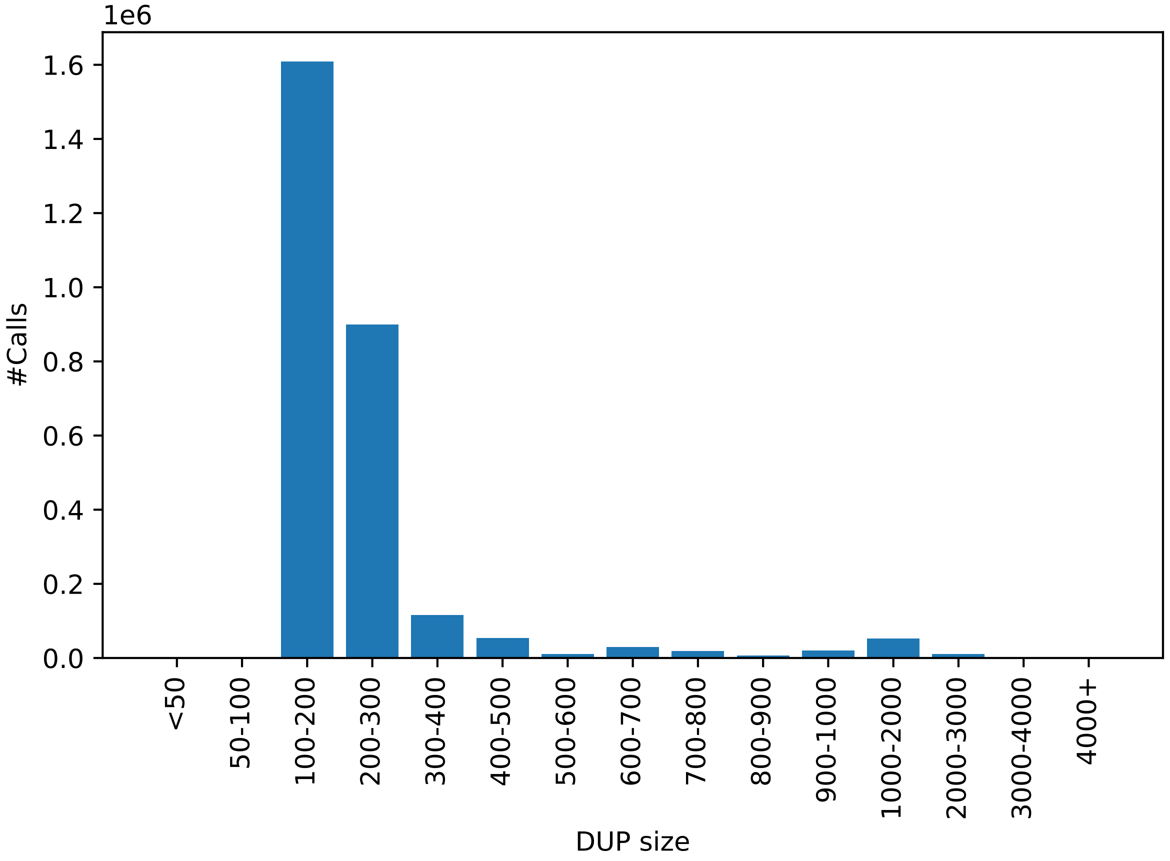


**Supplementary Figure 12.** Distribution of duplication call (merged CNV segments) sizes in CNVnator predictions on the (WGS) 1000 Genomes data set (training set).

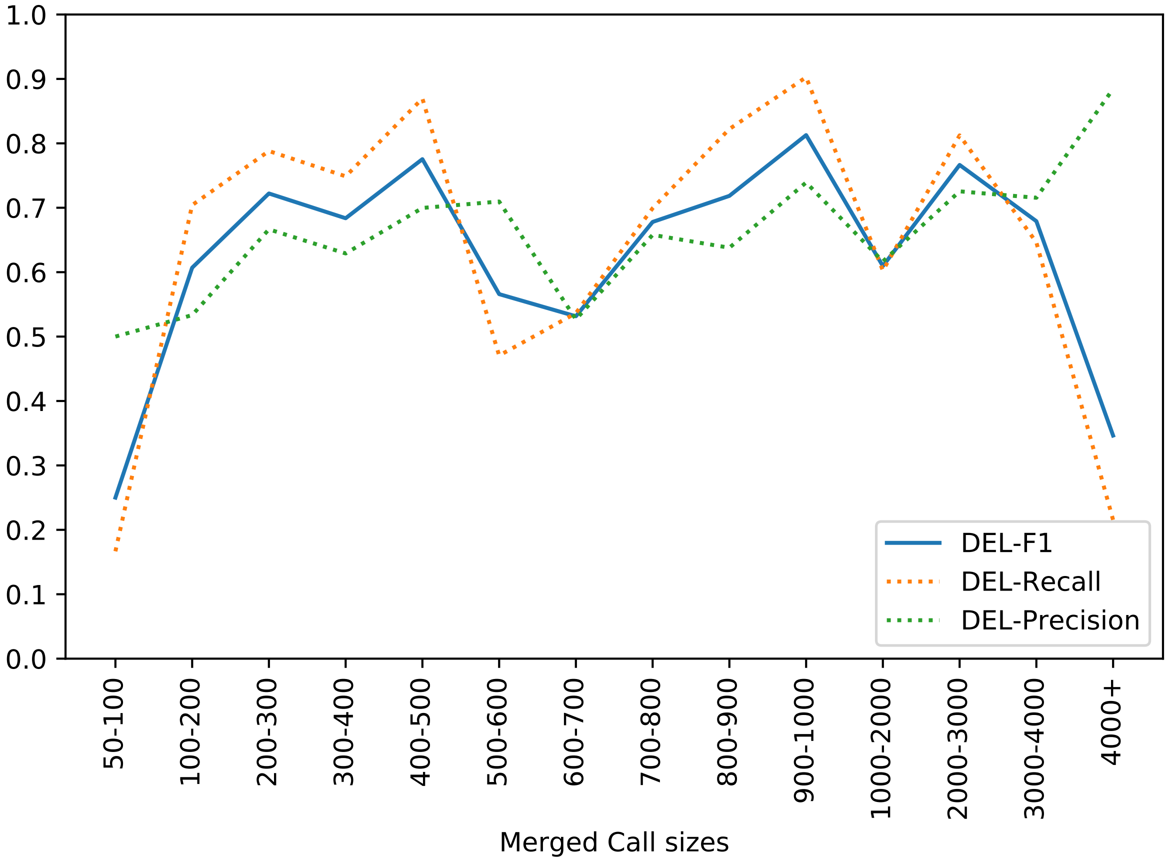


**Supplementary Figure 13.** Deletion detection performance (F1 score, Recall, Precision) of ECOLE across different call sizes for merged calls.

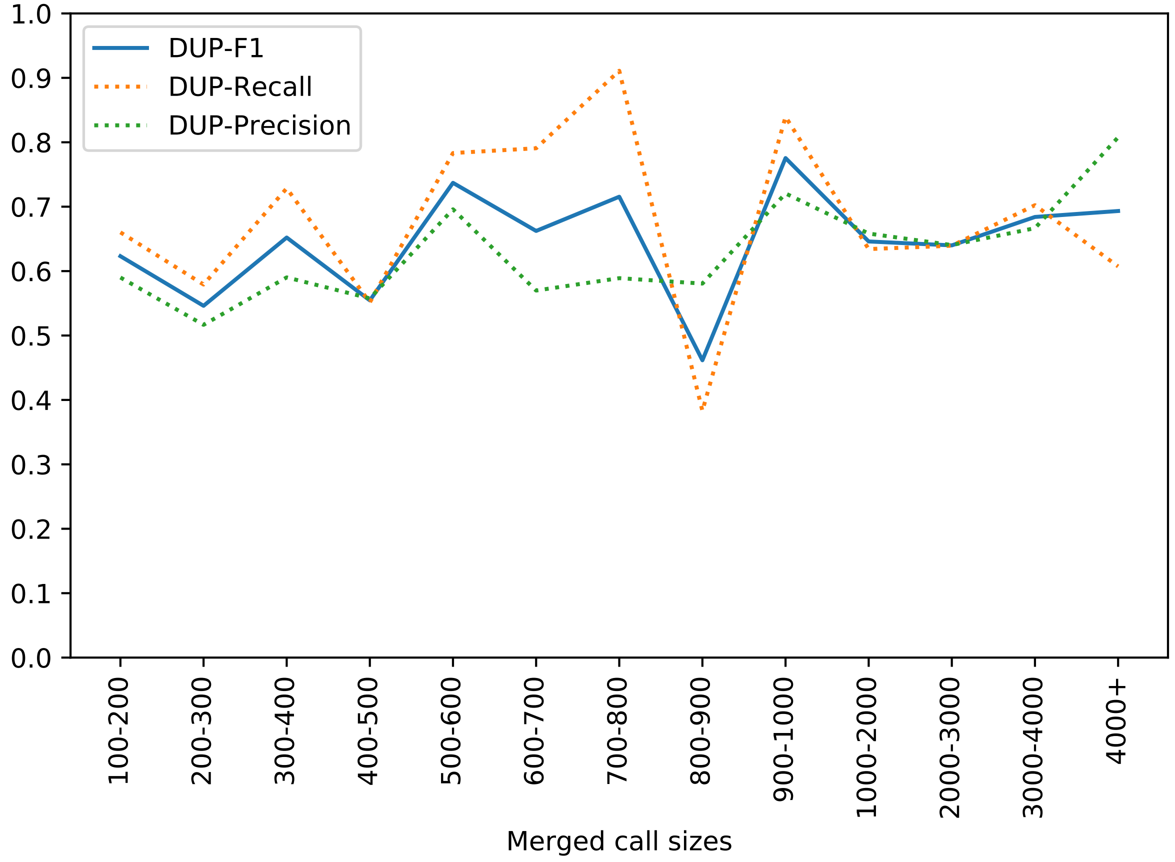


**Supplementary Figure 14.** Duplication detection performance (F1 score, Recall, Precision) of ECOLE across different call sizes for merged calls.

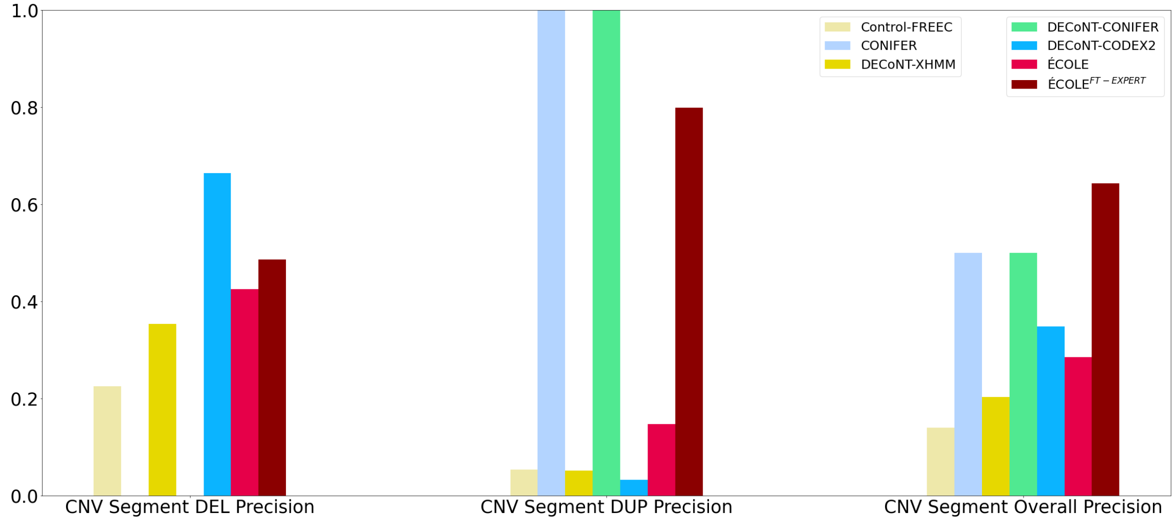


**Supplementary Figure 15.** The performance comparison of the WES-based CNV callers on the Chaisson et. al. data set (test set) on CNV segments. CNVnator calls on the matched WGS samples are used as the ground truth. Control-FREEC returns exact (integer) copy number predictions, which are discretized into deletion, duplication, and no-call. We also used the DECoNT tool to polish call sets of all considered tools which are denoted by DECONT-*tool_name*

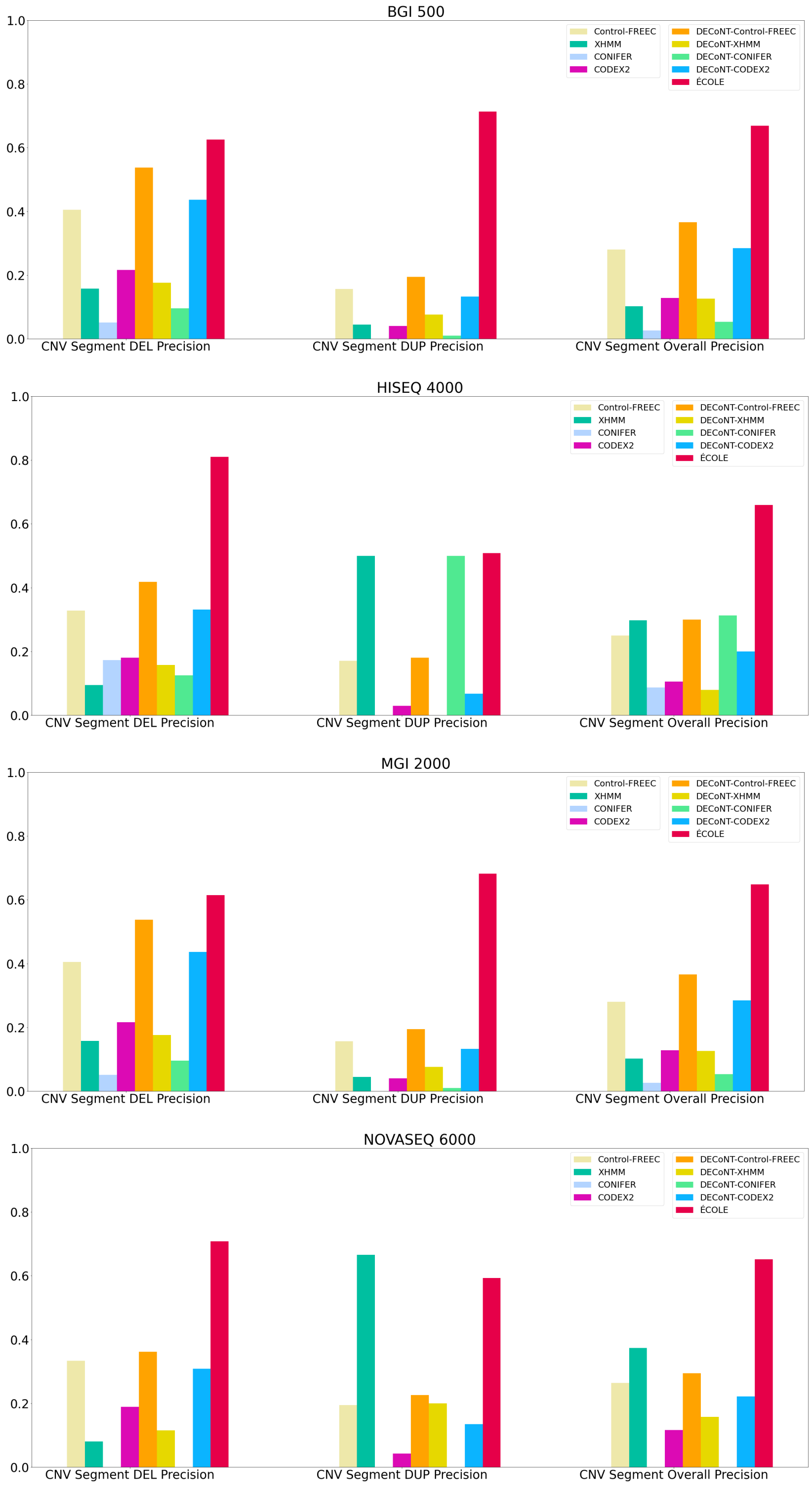


**Supplementary Figure 16.** Performance comparison with other tools on NA12878 sample on CNV segments. See Supplementary Table 23 for the values used to produce this plot and see Supplementary Tables 24-27 for the corresponding confusion matrices.

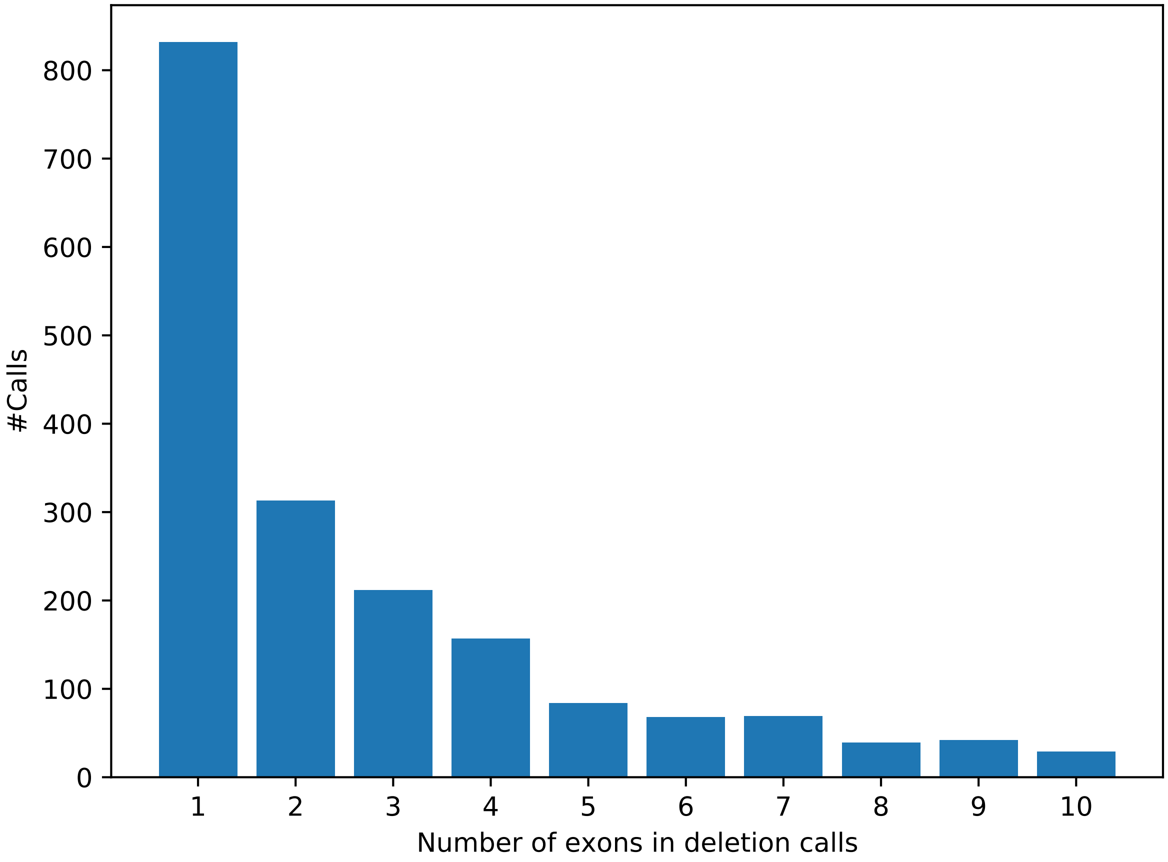


**Supplementary Figure 17.** Distribution of the number of exons in the merged deletion calls.

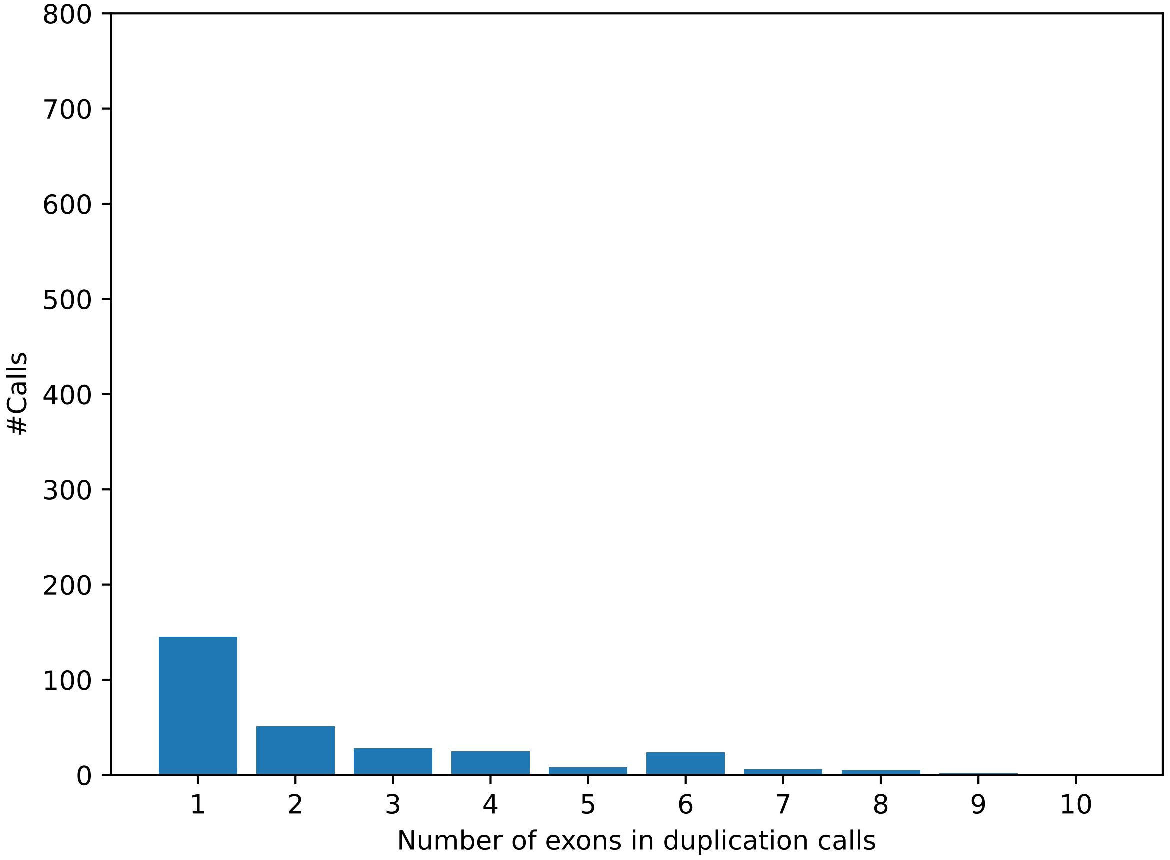


**Supplementary Figure 18.** Distribution of the number of exons in the merged duplication calls.

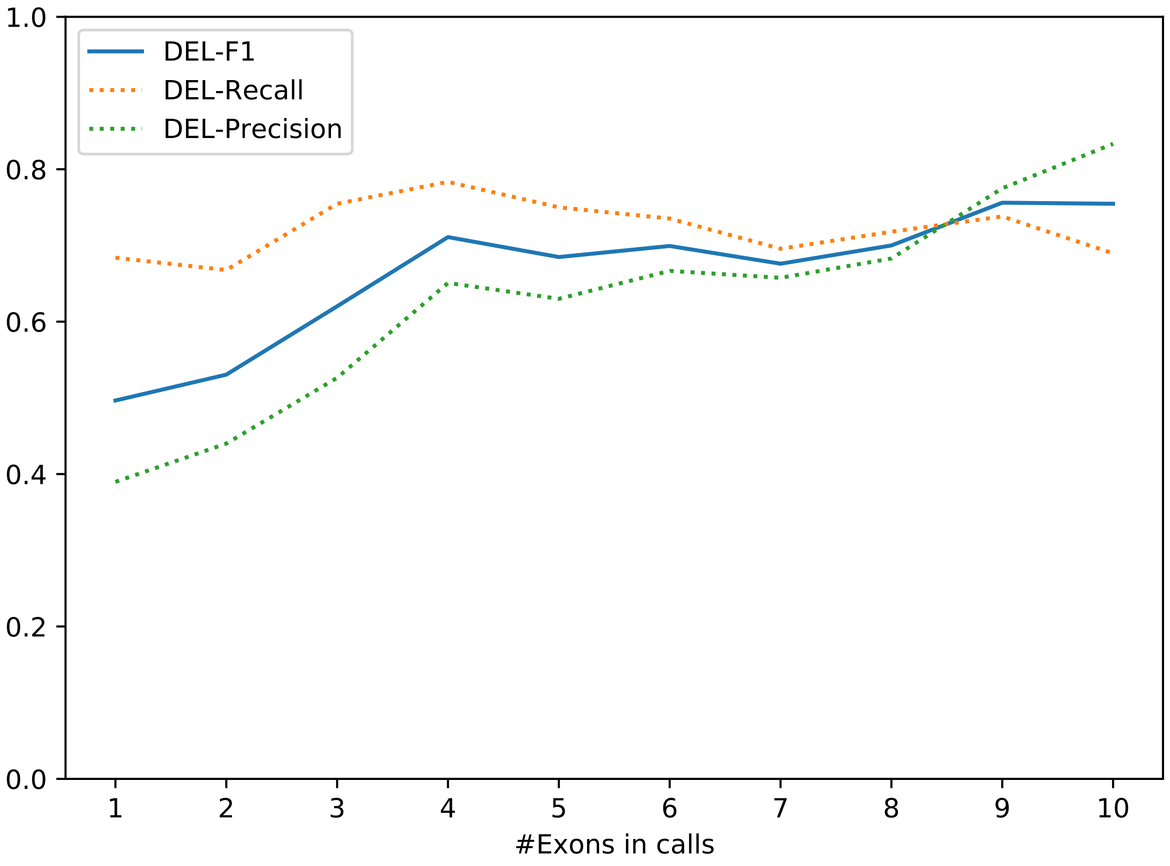


**Supplementary Figure 19.** Deletion performance (F1 score, Recall, Precision) of ECOLE by the number of exons in calls.

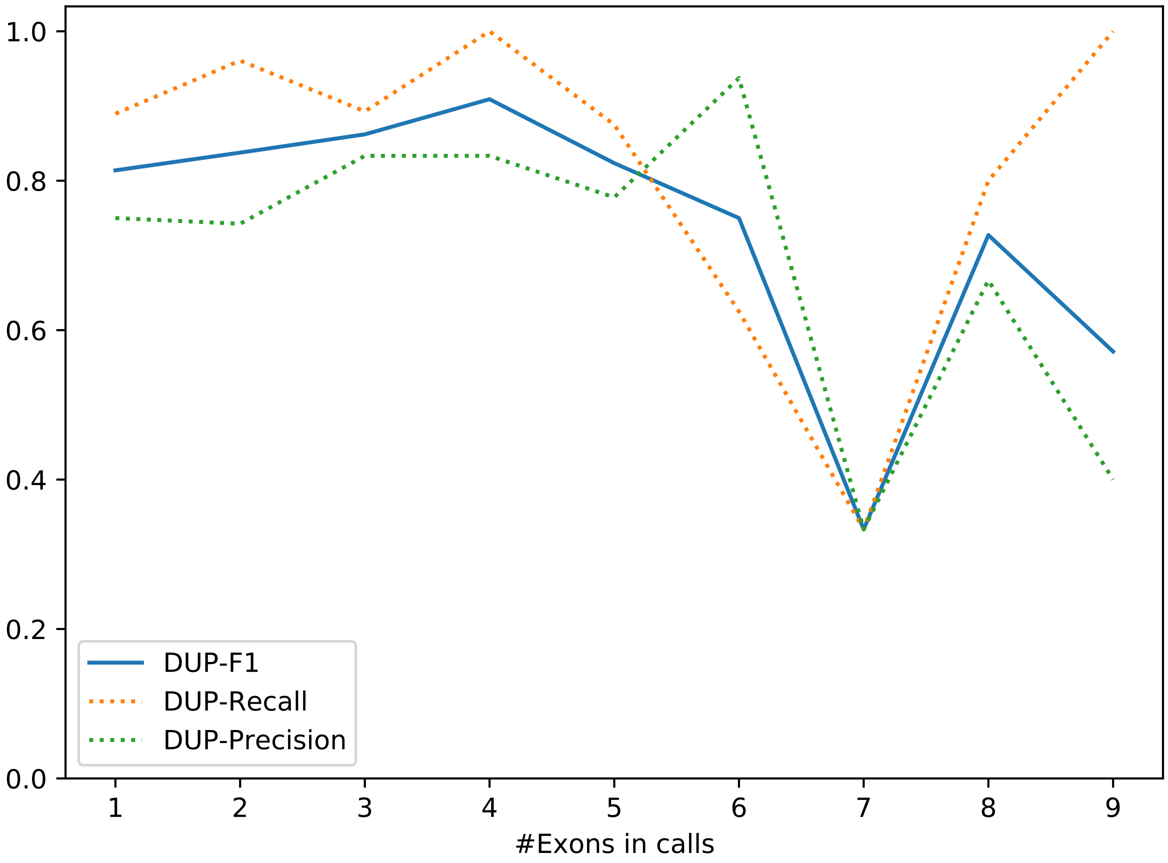


**Supplementary Figure 20.** Duplication performance (F1 score, Recall, Precision) of ECOLE by the number of exons in calls.

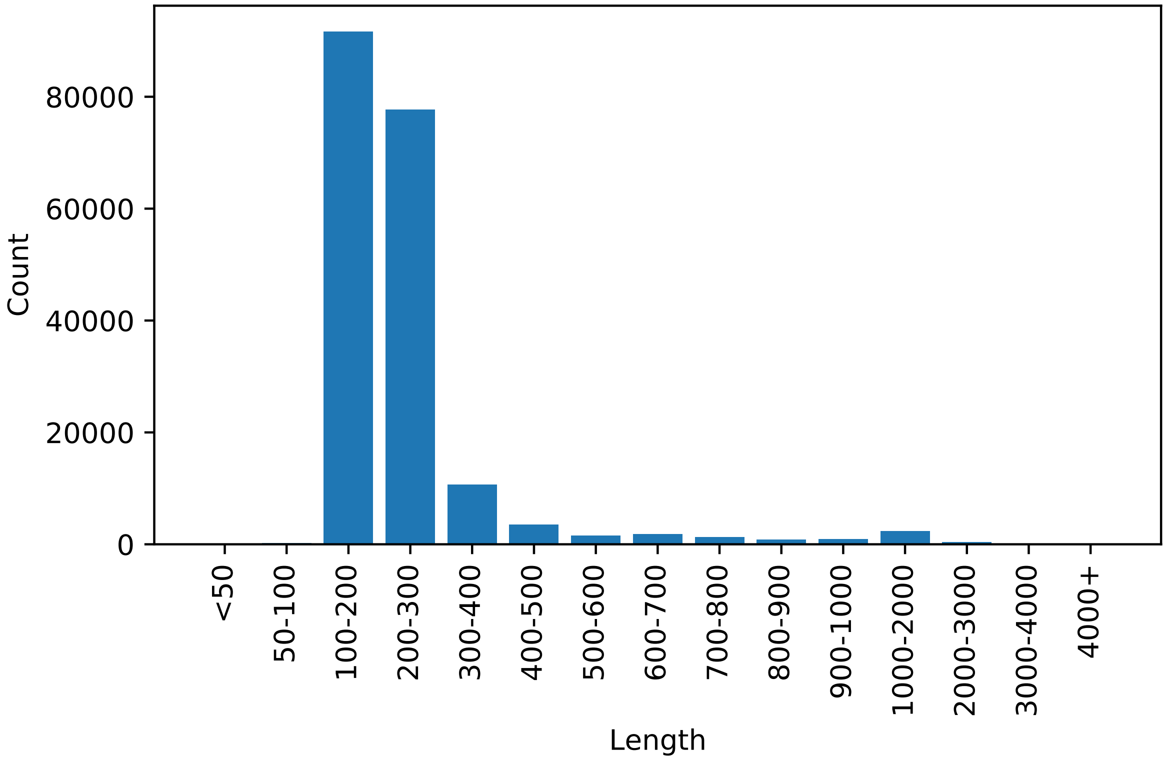


**Supplementary Figure 21.** The length distribution of exons

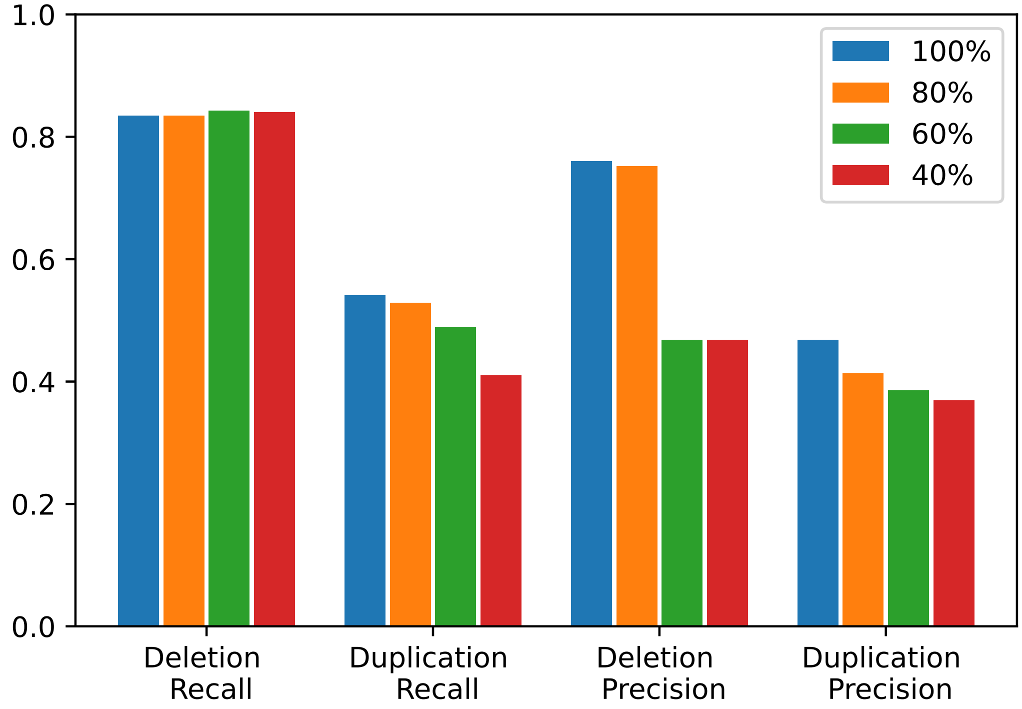


**Supplementary Figure 22.** The performance of ECOLE$^{\text{FT-GIAB}}$ on NA12892 samples subsampled at the rate of 80%, 60%, and 40%.

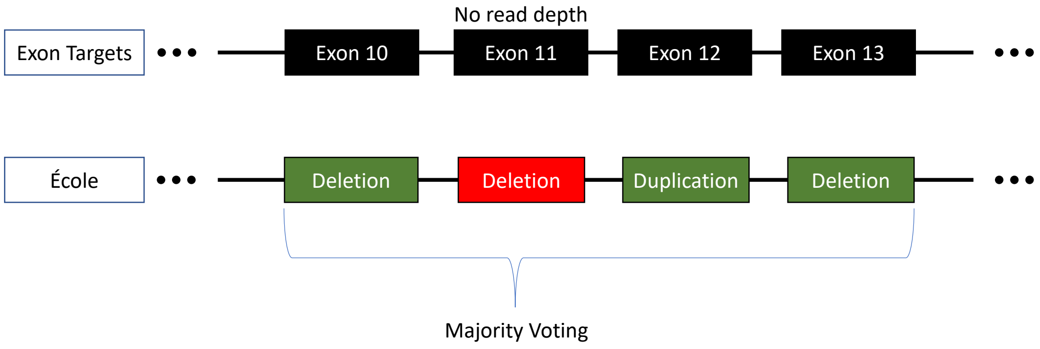


**Supplementary Figure 23.** Some exons do not have read depth information, thus ECOLE cannot make CNV calls for these exons immediately. Initially, ECOLE calls are obtained for exons that have read depth value. Later, the calls for exons that don't contain read depth information are obtained by applying majority voting on a window of size 3 closest exons on the ECOLE calls for neighboring exon regions. In this example, the red exon did not have a read depth signal and inferred using this method.

### 3 Supplementary Notes

#### 3.1 Derivation of relevance scores for ECOLE interpretability

The attention mechanism introduced in [5] is as follows:

$$\mathbf{A} = softmax(\mathbf{Q} \cdot \mathbf{K}^{)} \quad (1)$$
$$\mathbf{O} = \mathbf{A} \cdot \mathbf{V} \quad (2)$$

where $\mathbf{O} \in R^{h \times s \times d_h}$ is the output of the multi-head attention, $\mathbf{Q} \in R^{h \times s \times d_h}$ is the query matrix, $\mathbf{K}, \mathbf{V} \in R^{h \times s \times d_h}$ are the key and value matrices. $h$ is the number of heads in multi-head attention module, $d_h$ is the size of the embedding dimension. $\mathbf{A}$ inherently defines the connections between each token along the sequence of length $s$. In order to fetch the relevance of the attention signals with respect to the classification, we will use the relevance scores defined in [2]. The relevancy scores define the importance of each token for the predicted class of the input. We initialize the relevance scores with an identity matrix as follows:

$$\mathbf{R} = I_s \quad (3)$$

where the identity matrix is $s \times s$ and $s$ denotes sequence length. This explainability method uses the attention $\mathbf{A}$ matrix from each attention head and uses the gradient signals to average across the heads. The resulting attention map $\bar{\mathbf{A}} \in R^{s \times s}$, is as follows:

$$\bar{\mathbf{A}} = E_h((\nabla \mathbf{A} \odot \mathbf{A})^+) \quad (4)$$

where $\odot$ is the Hadamard product between the gradient matrix and the attention matrix $\nabla \mathbf{A} := \dfrac{\partial y_t}{\partial \mathbf{A}}$, and $y_c$ is the prediction of the model for the class $c$ we perform explanation, and $E_h$ is average over the multiple attention heads.

$$\mathbf{R} = \mathbf{R} + \bar{\mathbf{A}} \cdot \mathbf{R} \quad (5)$$

We use the Attention matrices $\mathbf{A}$ of the last Transformer layer and obtain the specified relevance scores. Moreover, we are using Performer [3] with FAVOR+ mechanism which approximates self-attention and calculates the output without using the attention matrix explicitly. Therefore, we reconstruct the attention matrix explicitly by replacing the value matrix in Equation 2, with an identity matrix that contains the indices of the sequence tokens.

#### 3.2 Hyperparameter Tuning for Baseline Models

First, we consider linear SVM classifier and XGBoost with the default hyperparameters. We trained these 2 baseline methods on our training set (1000 Genomes) for 150 rounds (partial fit). We then tested the trained methods using the 1000 Genomes test data set. The performance benchmark of these methods with default parameters and ECOLE can be seen in Supplementary Table 8.

To make these methods stronger baselines, we decide to tune the hyperparameters. For this, we adopt a 5-fold cross-validation routine and applied a grid search on the validation set to select the best hyperparameters for the linear SVM classifier and XGBoost. We selected the validation set for each batch of the training set in the current round. We set the validation set as 20% of the training set. We used a random seed for this selection. We then trained these 2 methods using the best hyperparameter setting for each batch and round of the training. Finally, we tested the trained methods with tuned hyperparameters. However, the SVM classifier and XGBoost with tuned hyperparameters performed similarly to the methods with default hyperparameters, Thus, we chose to include the default SVM classifier and XGBoost as our baseline model.

The hyperparameter search space for linear SVM classifier was as follows: (1) the loss function to be used $\in \{'hinge','log_loss','squared_error'\}$; (2) the penalty to be used $\in \{"l2","l1","elasticnet", None\}$; (3) constant that multiplies the regularization term sampled 5 equidistant learning rates from $[10^{-3}, 10^{-4}]$ and (4) the maximum number of passes over the training data $\in \{500, 1000, 2000\}$. The hyperparameter search space for XGBoost was as follows: (1) the loss function to be used $\in \{`log_loss', `exponential'\}$; (2) the function to measure the quality of a split. $\in \{`friedman_mse', `squared_error'\}$; (3) the minimum number of samples required to split an internal node $\in \{2, 5, 10\}$; (4) the minimum number of samples required to be at a leaf node $\in \{1, 2, 5\}$ and (5) maximum depth of the individual regression estimators $\in \{2, 3, 4\}$.

##### 3.3 Chromosome-specific CNN model

This CNN-based model contains 4 layers of 1-dimensional convolutional neural network and 1 linear layer of 311 neurons. The kernel sizes of the 4 layers of CNN are as follows: 100, 100, 160, 160. We use ReLU as activation function. We use Max Pooling and Dropout between the last CNN and linear layer. We trained 2 different CNN models for chromosomes 21 and 10. In Table 31, the results of these 2 chromosome-specific CNN models along with ECOLE are shown. Note that we trained and tested these 2 chromosome-specific CNN-based models on the same respective chromosomes. On chromosome 21, even though the CNN-based model attains a high precision (58.8%), ECOLE is able to outperform this model in both precision (85.5%) and recall (92.6%). On chromosome 10, the CNN-based model performs very poorly and attains 0% precision and recall. On the other hand, ECOLE performs better than the CNN-based solution and attains 3.7% precision and 29.2% recall. We observe that CNN-based models perform much worse than ECOLE in their respective chromosomes.

##### 3.4 The Architecture of Generic CNN model

The CNN model contains 5 layers in total - 4 convolutional layers and a fully connected layer. The first CNN layer is a 1D convolutional layer containing 100 kernels with a size of 10 with stride 1. It takes the input data $R^{1024}$, returns an output vector $\in R^{100x1015}$ followed by ReLU activation. The second convolutional layer uses 100 kernels with size 10. It generates the vector $\in R^{100x1006}$ again followed by ReLU activation and max pooling to, resulting in the output $\in R^{100x335}$. Subsequently, two more 1D convolutional layers are applied with kernel sizes of 160, and ReLU activation is performed after each convolutional layer. It returns a vector $\in R^{160x317}$. Then, global max-pooling is applied resulting in a 317 dimensional vector. To prevent overfitting, a dropout layer is used. Finally, a fully connected layer is used to map the output of the previous layer to the desired 3-dimensional output followed by softmax activation. Each dimension corresponds to labels: DEL, DUP, and no-call, respectively. We use the weighted cross-entropy loss function.
